# The RNA landscape of the human commensal *Segatella copri* reveals a small RNA essential for gut colonization

**DOI:** 10.1101/2024.04.05.588293

**Authors:** Youssef El Mouali, Caroline Tawk, Kun D. Huang, Lena Amend, Till Robin Lesker, Falk Ponath, Jörg Vogel, Till Strowig

## Abstract

The bacterium *Segatella copri* is a prevalent member of the human gut microbiota associated with both health and disease states, but intrinsic factors that determine its ability to effectively colonize the gut are not understood. By extensive transcriptome mapping of *S. copri* and examining human-derived samples, we discover a previously unknown small RNA named here *Segatella* RNA colonization factor (SrcF). We show that SrcF is essential for *S. copri* gut colonization in a gnotobiotic mouse model. SrcF regulates genes involved in nutrient acquisition, and its own expression is controlled by complex carbohydrates, particularly fructans. Furthermore, SrcF expression is strongly influenced by human microbiome composition and by the breakdown of fructans by cohabitating commensals, suggesting that the breakdown of complex carbohydrates mediates inter-species signaling among commensals beyond its established function in generating energy. Together, this study highlights the contribution of a small RNA as a key regulator in gut colonization.

## INTRODUCTION

Human microbiomes represent complex microbial ecosystems comprised of thousands of species, many of which are specialized to their organ site or even to the human host. Within these ecosystems, microbes compete or cooperate to acquire essential nutrients, in particular, carbohydrates. Work in the model gut species *Bacteroides thetaiotaomicron* has discovered sophisticated networks of transcriptional and posttranscriptional control that help adjust bacterial gene expression in response to carbon availability ^1–3^. However, how other prevalent non-model gut commensals sense and prioritize nutrient acquisition for successful gut colonization remains unexplored. This is particularly true for *Segatella copri* (formerly known as *Prevotella copri*) ^4^, which is a prevalent and exceptionally abundant member (most abundant in 34% of individuals) ^5,6^ of the human gut microbiota where it constitutes a keystone species dominating one of the three proposed human enterotypes ^7^.

*S. copri* is part of the *S. copri* complex containing 13 different species, which are ubiquitous in traditional agrarian, non-Westernized societies that are known to consume fiber-rich diets ^8–10^. Yet, the *S. copri* complex is less prevalent in Westernized societies ^5,11^, where its presence has been disparately associated with inflammatory diseases ^12–16^ or health-beneficial effects ^17^. On the former, the presence of *S. copri* seems to correlate with the onset of rheumatoid arthritis in humans ^12,15^ and certain *S. copri* isolates exacerbate this disease in a mouse model ^16^. On the latter, the presence of *Segatella* spp. is positively correlated with a vegetarian or a vegan diet, both of which are rich in complex polysaccharides ^18,19^. In agreement, *S. copri* isolates can readily utilize a wealth of plant-derived polysaccharides ^5,20,21^. This nutritional versatility results from a multiplicity of gene clusters named polysaccharide utilization loci (PUL), which encode both regulators and enzymes for the breakdown and transport of polysaccharides and their degradation products ^2,22^.

*S. copri* encodes for a few dozen PULs specialized for the utilization of a defined carbon source, some of which have been experimentally validated ^21^. By contrast, it is less clear how *S. copri* senses and prioritizes nutrient acquisition for successful gut colonization. While metagenomics-based studies have delved into *S. copri* complex genomic content ^5,19,20^, we lack any knowledge on transcriptional and/or translational mechanisms in *S. copri,* as to which promoter sequences drive gene expression or how translation of mRNAs is regulated. Consequently, the post-transcriptional RNA landscape remains unexplored as to whether small non-coding RNAs are present in *S. copri*, and how they might contribute to integrating environmental cues in the gut to impact gene expression and successful colonization.

To better understand how *S. copri* controls gene expression for successful gut colonization, we have generated a comprehensive RNA map of *S. copri in vitro* and *in vivo* that uncovers fundamental aspects of gene regulation, including the presence of sRNAs with potential regulatory functions. To identify regulators of gut colonization, we interrogated human-derived samples and identified a highly expressed sRNA in human microbiomes that is conserved in *S. copri* strains, and showed it is essential for gut colonization in gnotobiotic mice. We named it SrcF, for *Segatella* RNA colonization factor, and uncovered that its deletion strongly impacts *S. copri* gene expression, including that of several PULs. Furthermore, we have pinpointed dietary components, such as fructans, as environmental cues dictating SrcF expression. This expression modulation occurs through the release of monosaccharides from *S. copri*-accessible fructans, as well as from inaccessible fructans degraded by cohabitating microbiota members indicating that extracellular breakdown of complex carbohydrates may play a role in inter-species signaling beyond its attributed function as solely an energy source. Importantly, our results highlight the relevance of non-coding sRNAs in the biology of a prevalent gut commensal and underscore the necessity to extend beyond the bacterial coding genome to fully capture commensal interactions in the human gut microbiome.

## RESULTS

### The primary transcriptome of *Segatella copri* HDD04 and DSM18205^T^

The genome-wide RNA landscape of *Segatella copri* was determined by applying differential RNA-seq (dRNA-seq) that aims to globally identify the 5’ends of transcripts by distinguishing primary (5’-PPP) transcripts ends from processed (5’-P, 5’-OH) transcripts via a specific nuclease treatment ^23^. dRNA-seq assigns transcriptional start sites (TSSs) to annotated coding sequences (see methods) and identifies non-coding unannotated genetic elements with potential regulatory function. Here, we subjected *S. copri* HDD04 and DSM18205^T^ strains, representing a genetically tractable isolate and the type strain ^21,24^, respectively, to dRNA-seq at three growth phase conditions namely early exponential phase (EEP; OD_600nm_ 0.25; ∼ 4h), mid-exponential phase (MEP; OD_600nm_ 0.8; ∼6h), and stationary phase (Stat; OD_600nm_ 4.0; ∼11h) (Fig. 1A).

**Figure 1.**
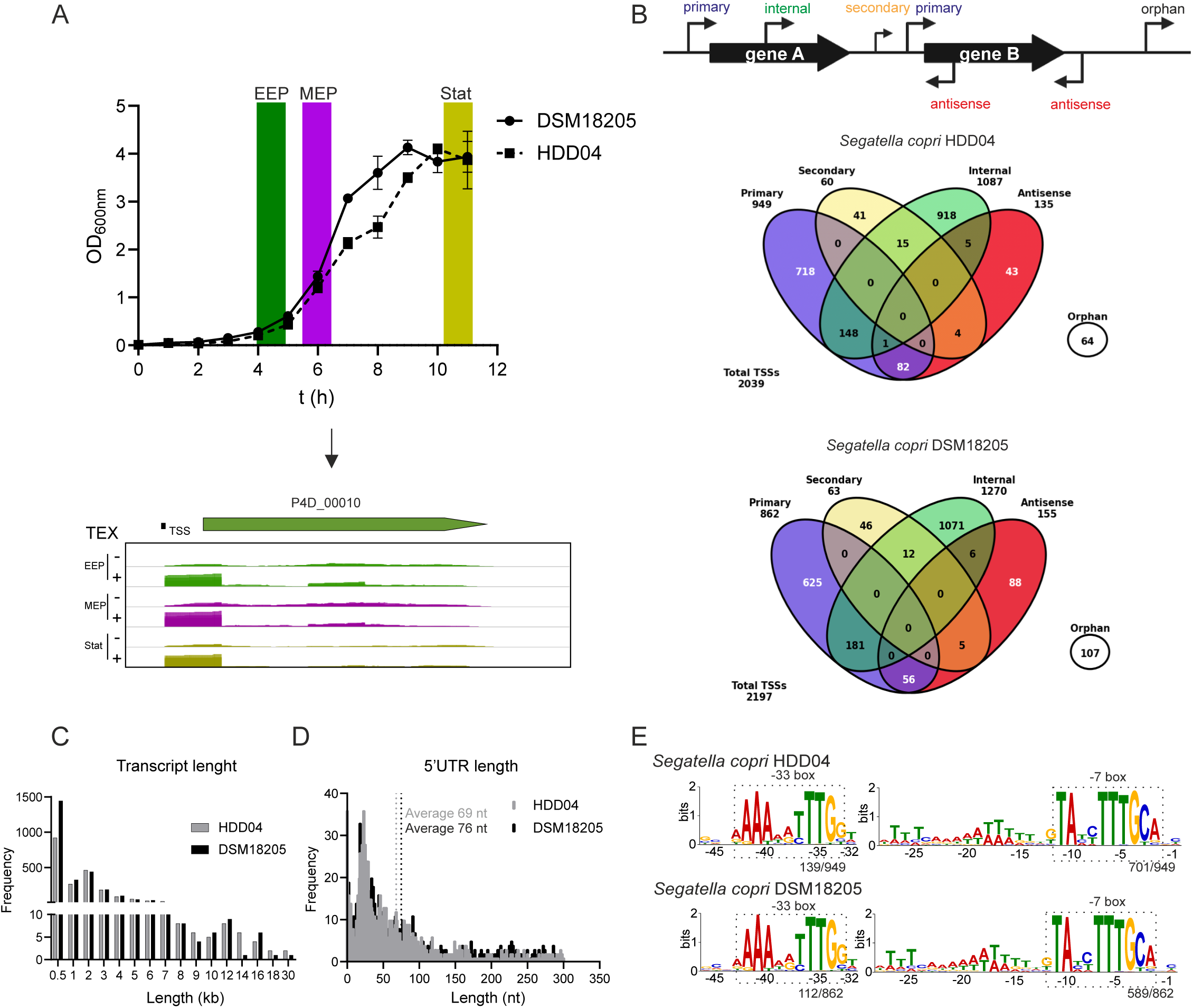
The primary transcriptome of *Segatella copri* HDD04 and DSM18205^T^. In the upper panel, growth curves of *S. copri* HDD04 and DSM18205^T^ in BHI-S. Highlighted time-points indicate the OD_600nm_ at which samples were collected for dRNA-seq at the early exponential phase (EEP), mid-exponential phase (MEP), and stationary phase (Stat). Lower panel, an IGV Genome Browser display of an example of TSS mapping of the *S. copri* HDD04 locus P4D_00010 with dRNA-seq. Indicated +/- for library subjected or not to Terminator Exonuclease treatment (TEX) treatment. **B.** Upper panel, schematic representation of TSS classification based on the location relative to the associated CDS. Lower panels, Venn diagrams showing the number of the 5 TSS categories detected in *S. copri* HDD04 and *S. copri* DSM18205^T^ by TSSpredator in +/- TEX libraries at EEP, MEP, and Stat. **C.** Representation of the number of transcripts with different transcript lengths detected by dRNA-seq analysis. Data for both *S. copri* HDD04 and DSM18205^T^ is shown. **D.** Representation of the frequency and length of 5’UTRs detected in *S. copri* HDD04 and DSM18205^T^ by dRNA-seq analysis. **E.** Promoter motif detected 50 nt upstream of primary TSSs in *S. copri* HDD04 and DSM18205^T^. Promoter motifs were identified by MEME and the frequency of the motifs is indicated for the -7 box and -33 box.

Genome-wide identification of TSSs using ANNOgesic ^25^ led to the identification of over 2,000 TSSs in both *S. copri* HDD04 and DSM18205^T^ that were classified into five categories, with a similar distribution in both strains, based on their genomic location in relation to the associated coding sequence (CDS) (Fig. 1B, see methods). Predicted TSSs that do not associate with any CDS were classified as orphan and might be indicative of intergenic unannotated genetic elements (see below) (Supplementary Table S1). The HDD04 and DSM18205^T^ strains display similar transcript length derived from TSS and terminator predictions (Fig. 1C, see methods). Moreover, the identification of the 5’ untranslated regions (5’UTR) of mRNAs from primary TSSs and CDS shows a similar average length in *S. copri* HDD04 and DSM18205^T^ 5’UTRs with 69 nt and 76 nt, respectively (Fig. 1D). In addition, a similar number, 36 and 35 respectively, of leaderless transcripts (TSS at start codon) are identified in both strains (Fig. 1D). As does *Bacteroides thetaiotaomicron* ^26^, *S. copri* lacks a clear consensus ribosome binding site.

To identify promoter sequences, the 50 nt sequence regions upstream of the annotated primary TSSs were analyzed for consensus sequences ^27^. Notably, a highly conserved -7 box TAnnTTTCA was identified in over 70% of primary TSS in HDD04, and a conserved -33 box AAAnnTTTGG was identified only in 15% of the primary TSSs of *S. copri* HDD04 (Fig. 1E). Identical promoter motifs were found in *S. copri* DSM18205^T^ with a similar prevalence (Fig. 1E). The promoter motifs identified here resemble the canonical promoter motif recently identified in *B. thetaiotaomicron* ^26^. Together, these results support a conserved basic transcriptional machinery among Bacteroidota.

### Non-coding RNA repertoire in *Segatella copri* and conservation across the *Segatella* species complex

The annotation of TSSs of the primary transcriptome of *S. copri* via dRNA-seq allows the mapping of non-annotated regulatory RNA elements including non-coding small RNAs (see methods). First, we validated the expression of housekeeping RNAs annotated via Infernal ^28^ in *S. copri* HDD04 and DSM18205^T^. Both strains encode for transfer-messenger RNA (tmRNA) and the RNA component of RNase P (M1 RNA) (Fig. 2A, S1A). In addition, they encode for Bacteroidales-1 RNA (RFAM, RF01693) ^29^, recently identified as a homolog of the 6S RNA in *B. thetaiotaomicron* ^26^. The expression of housekeeping RNAs was validated via northern blotting for the three selected growth conditions (Fig. 2A, S1A) and a similar profile of expression was detected in both strains with all three RNAs accumulating at the stationary phase, the latter, a characteristic feature of the 6S RNA ^30^ (Fig. 2A, S1A). Given the high similarity in *S. copri* HDD04 and DSM18205^T^ primary transcriptome and given the over 95% average nucleotide identity of their genomes ^31^, further characterization of the primary transcriptome was carried out in genetically tractable *S. copri* HDD04.

**Figure 2.**
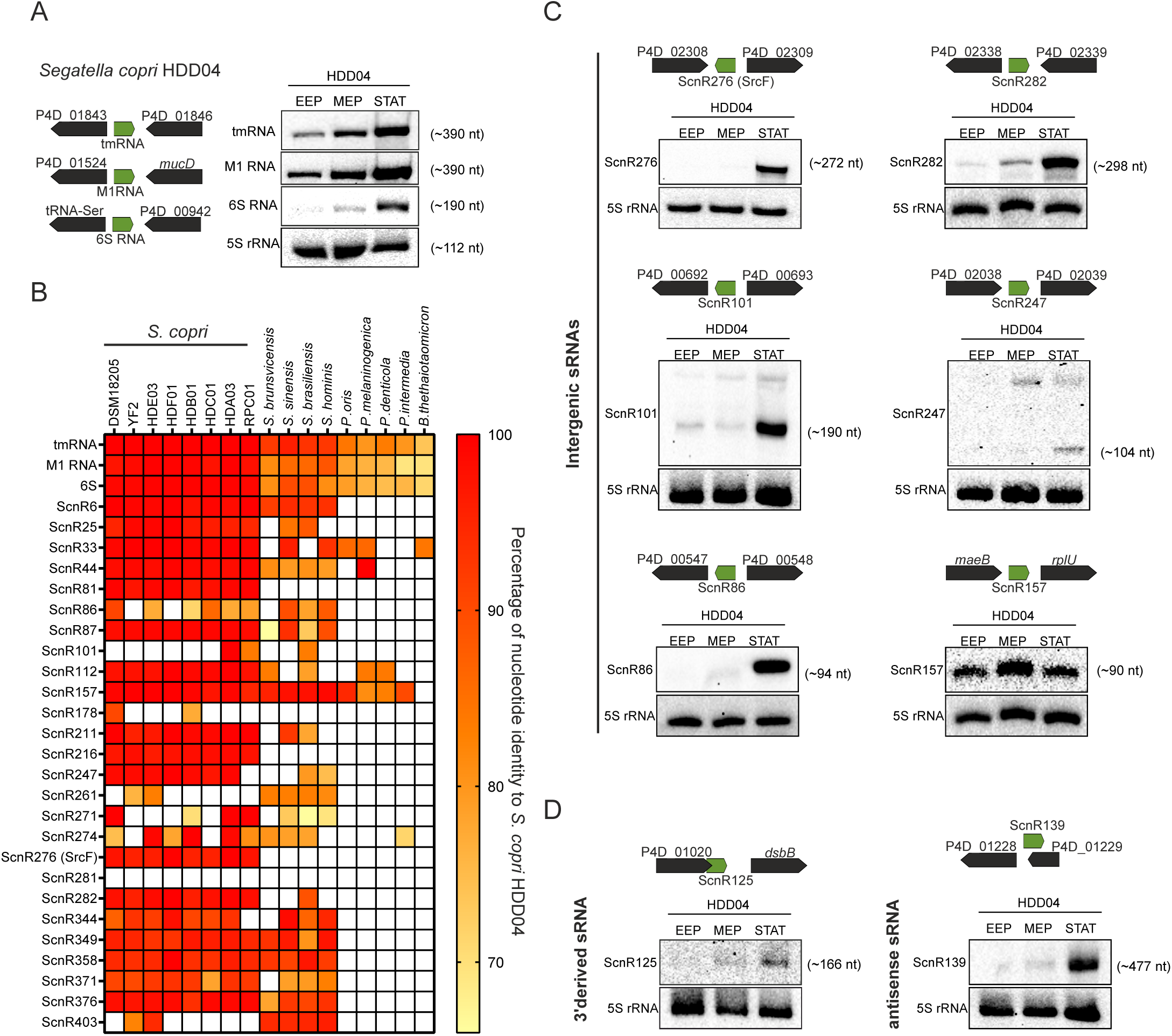
Non-coding RNA repertoire in *Segatella copri* and conservation across the *Segatella* species complex. Northern blot detection of housekeeping non-coding RNAs tmRNA, M1RNA, and 6S RNA in *S. copri* HDD04. On the left, the genomic context of the corresponding RNAs is represented. On the right, detection of expression in total RNA samples at EEP, MEP, and Stat. 5S rRNA was detected as a loading control. **B.** Conservation analysis of housekeeping RNAs and the 27 intergenic non-coding sRNAs identified by dRNA-seq in *S. copri* HDD04. The sRNAs presence was determined by BLASTN and sRNAs with over 60% coverage and 70% identity were included. **C.** Northern blot detection of identified intergenic sRNAs: ScnR276 (SrcF), ScnR282, ScnR101, ScnR247, ScnR86, and ScnR157. ScnR: *Segatella copri* non-coding RNAs. 5S rRNA was detected as a loading control. **D.** Northern blot detection of ScnR125 and ScnR139 as examples of other sRNA categories. 5S rRNA was detected as a loading control.

The non-coding small RNAs (sRNAs) repertoire was comprehensively annotated combining ANNOgesic predictions and manual curation of the dRNA-seq data (see methods). In *S. copri* HDD04, a total of 63 non-coding RNAs were identified with an average length of 244 nt (range 75 to 477 nt) (Fig. S1B). Of those, 27 were encoded in intergenic regions, 7 were encoded in the antisense orientation to other genes, and the remaining 30 were either 3’ or 5’ derived from *S. copri* HDD04 mRNAs (Supplementary Table S1F). Of note, 6 of the intergenic sRNAs and 8 of the mRNA-derived sRNAs contain small open reading frames (< 50 amino acids) and might represent dual-function RNAs as previously described in other bacteria (Fig. S1B) ^32^. Conservation analysis of the intergenic sRNAs (BLASTN with 70% nucleotide identity) indicates that most of the sRNAs identified in *S. copri* HDD04 are conserved among *S. copri* (former *P. copri* clade A) strains (Fig. 2B). Over 60 % of the sRNAs are conserved in at least two additional isolates of *Segatella* spp, while in contrast, there is little conservation beyond the *Segatella copri* species complex, i.e., absent in other members of the Prevotellaceae family or *B. thetaiotaomicron* (Fig. 2B). Next, the expression of a subset of putative sRNAs in *S. copri* HDD04 was experimentally validated by northern blot detection. We detected six different intergenic sRNAs ranging from 90 to 300 nt at EEP, MEP, and STAT (Fig. 2C). Five of the intergenic sRNAs (ScnR282, ScnR276, ScnR86, ScnR101, and ScnR247) accumulated at stationary phase, while ScnR157 was constitutively expressed throughout the growth phases (Fig. 2C). Interestingly, ScnR157 is strongly conserved in *Segatella spp* (Fig. 2B), and its expression was experimentally validated in three additional species of the *Segatella* genus, namely *Segatella brunsvicensis*, *Segatella sinensis*, and *Segatella hominis* (Fig. S1C). While genomic synteny is only partially maintained (Fig. S1D), the sRNA sequence and the promoter region are highly conserved indicating the promoter regions identified in *S. copri* (Fig. 1E) seem to also drive gene expression in other *Segatella spp* (Fig. S1E). Lastly, the expression of sRNAs from other classes, ScnR125 (3’derived) and ScnR139 (antisense) was also detected (Fig. 2D), indicating that mRNA-derived sRNAs and antisense sRNAs may also be potential regulatory elements in *S. copri*.

### Primary transcriptome map of *S. copri* in the murine gut

The large intestine constitutes the primary habitat for *S. copri*, and despite the large-scale resolution of TSSs we obtained *in vitro*, we asked whether we could identify *in vivo* specific TSSs not expressed *in vitro*. For instance, while *S. copri* HDD04 encodes for 29 polysaccharide utilization loci (PULs) ^21^, 17 were low or not expressed *in vitro* preventing the mapping of the majority (∼60%) of PUL TSSs. While *S. copri* is unable to mono-colonize germ-free mice receiving a regular chow diet ^17^, it can stably colonize mice within synthetic minimal communities ^33^. A prominent gnotobiotic model carrying a simplified synthetic community is the Oligo-Mouse-Microbiota (OMM12) mouse model ^34^. We colonized OMM12 mice with the *S. copri* HDD04 strain (Fig. 3A) that reached a relative abundance of >10% at day 7 in the feces and ranged between 13-22% at day 14 in the gastrointestinal lumen (Fig. 3B). To obtain total RNA for the identification of TSSs *in vivo*, distal colon (DC) contents, i.e., where *S. copri* reached the highest abundance, were used for RNA extraction and subsequent dRNA-seq. The use of a defined community allowed us to obtain deep coverage of our *S. copri* strain of interest, and readily filter out reads cross-mapping to other community members in the mouse gut (Fig. 3C, see methods).

**Figure 3.**
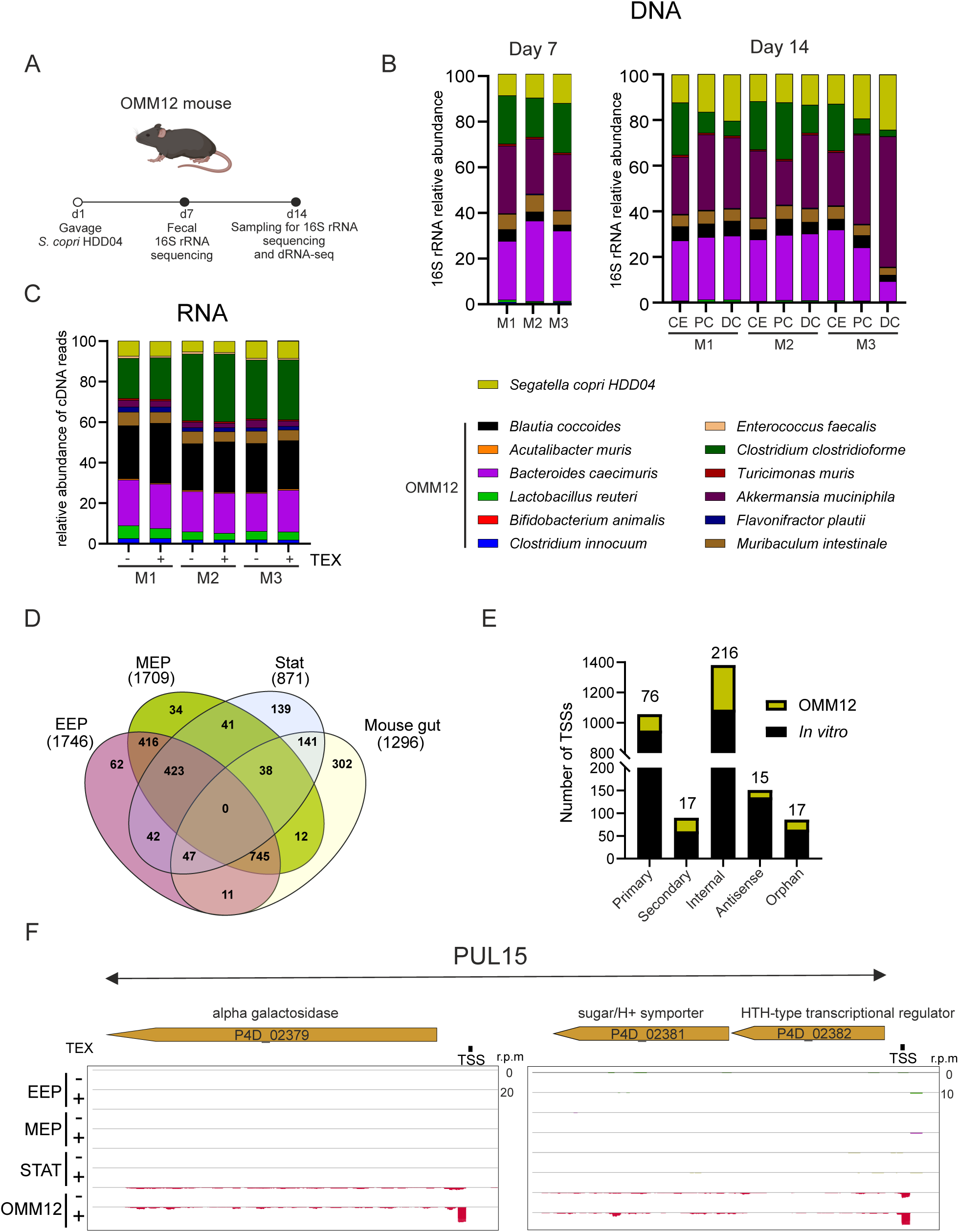
Primary transcriptome map of *Segatella copri* in the murine gut. Schematic of the workflow of OMM12 mouse model colonization with *S. copri* and sample collection. **B.** Relative abundance of *S. copri* HDD04 and the Oligo-Mouse-Microbiota (OMM12) commensals analyzed with 16S rRNA amplicon sequencing. Left panel detection from feces on day 7. Right panel, detection from the cecum (CE), proximal colon (PC), and distal colon (DC) at day 14 post gavage with *S. copri* HDD04. **C.** Relative abundance of reads mapped to *S. copri* HDD04 and OMM12 members in cDNA libraries generated from +/- TEX treated RNA samples. Samples were collected on day 14 from the distal colon of mice colonized with *S. copri*. **D.** Venn diagram of total TSSs detected in the four conditions tested, early exponential phase (EEP), mid-exponential phase (MEP), stationary phase (Stat), and the mouse gut. **E.** Number of total TSSs in each of the five categories. In yellow, the number of *S. copri* TSSs detected uniquely in the OMM12 mouse gut. **F.** Genomic visualization of the PUL15 locus TSS mapping in the four tested conditions. Indicated +/- for library subjected or not to Terminator Exonuclease treatment (TEX) treatment.

To identify *S. copri* TSSs in the mouse gut, reads aligned uniquely to *S. copri* were integrated into our dRNA-seq analysis, and transcriptional start sites (TSSs) were identified as described previously ^25^ (Supplementary Table S1B). TSSs detected in the mouse gut overlap to a large extent with TSSs detected in EEP and MEP samples rather than Stat samples (Fig. 3D). Interestingly, we detected an additional 302 TSSs (12% of total TSSs) unique to the OMM12 gut colonization (Fig. 3D-E). Of those, 76 represent primary TSSs (7% of total primary TSSs), associated with 11 of the 17 PULs in *S. copri* that were undetectable *in vitro*. For instance, we identified that the HTH-transcriptional regulator encoded by P4D_02382 in PUL15 is co-transcribed with a sugar/proton symporter (P4D_02381), whereas the expression of neither was detected *in vitro* (Fig. 3F). In addition to TSSs of PULs, we captured TSSs of additional genes such as P4D_631-633 that encode for Beta/alpha-amylases (Supplementary Table S1B) suggesting that *S. copri* could actively breakdown starch in the mouse gut. In agreement, genes within PUL15, also expressed in the gut, are predicted to break down starch ^21^. We did not detect the expression of 6 out of 29 PULs that putatively represent clusters of genes responsible for the breakdown of components not present in the mouse chow diet.

Overall, we generated a comprehensive map of *S. copri* transcriptional landscape both *in vitro* and *in vivo* that allowed the identification of over 1000 primary TSSs in *S. copri* (Fig. 3E). Interestingly, among newly identified TSSs, we did not annotate any additional sRNA indicating that no potential sRNA regulator is exclusively expressed in the mouse gut. However, the interrogation of the generated *S. copri* primary transcriptome under relevant environmental cues for *S. copri*, namely nutrient availability, contact with other commensals, or even directly within human gut microbiome communities will shed light on *S. copri* requirements for successful gut colonization.

### *Segatella* conserved sRNA SrcF is highly abundant in human gut microbiomes

Among the newly annotated elements, we identified intergenic non-coding RNAs that represent important posttranscriptional regulators in bacteria and are involved in fine-tuning gene expression ^2,37,38^. The deconvolution of complex ecosystems such as the human gut microbiome into defined environmental stimuli can be challenging. Nonetheless, we aimed to elucidate the contribution of sRNAs to *S. copri* colonization of the gut and explored a unique resource of isolated strains (*S. copri* HDA03, HDB01, HDC01, HDD04) for which we have obtained the metagenomes and metatranscriptomes from their respective donors (donors A, B, C, and D) (Fig. 4A-B). This allowed us to quantify the expression of the conserved sRNAs in human stool samples compared to when the *S. copri* isolates were grown *in vitro* (Fig. 4C) (Supplementary Table S2). Interestingly, ScnR282 and ScnR157 were among the most downregulated sRNAs in all donors (Fig. 4C). Similarly, in all donors we observe upregulation of ScnR276, while ScnR247 is upregulated in donors A, B, and D but not C (Fig. 4C). Analysis of the ScnR276 sequence, renamed here SrcF for *Segatella* RNA colonization factor (see below), shows high conservation at the nucleotide level and genomic synteny in all four donor strains (Fig. 4D-E).

**Figure 4.**
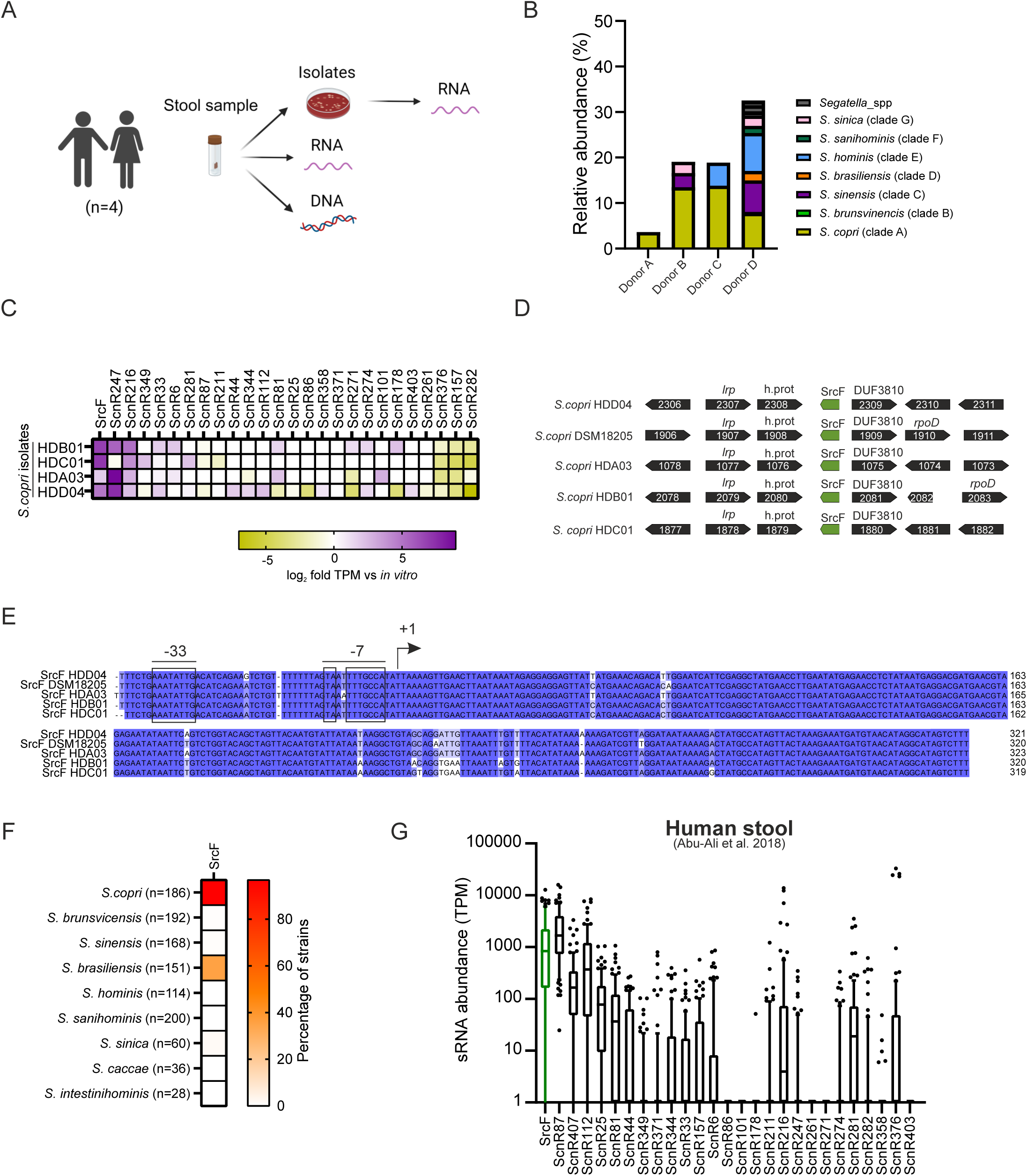
*Segatella* essential RNA colonization factor SrcF is highly abundant in human gut microbiomes. Schematic depiction of strain isolation and sample processing in 4 healthy human donors. **B.** Relative abundance of *Segatella spp*. in the four human donors determined by metagenome analysis. **C.** *S. copri* sRNA expression in human microbiomes. Heatmap representing log_2_ fold-change in *S. copri* sRNA expression within each donor in B compared to expression *in vitro*. **D.** Genomic context of SrcF loci in different *S. copri* isolates. **E.** Alignment of SrcF sequences from different *S. copri* strains including 50 nt upstream of the TSS. **F.** Heatmap representing the percentage of strains that encode for SrcF sRNA in the nine human *Segatella spp*. (n) represents the number of genomes analyzed in each species. Cut-off set to over 70% sequence identity and 60% coverage. **G.** Expression of sRNAs in human microbiomes carrying *S. copri*. The median of sRNATranscripts per Million (TPM) is represented. Samples with over 5% reads mapped to *S. copri* and with at least 5 sRNAs detected are included (n=82). Raw data in this panel obtained from Abu-Ali et al, 2018.

To interrogate human microbiomes carrying uncharacterized *S. copri* isolates, we extended the conservation analysis of the identified *S. copri* sRNAs beyond the selected isolates above and performed a systematic conservation analysis in all available sequenced *Segatella* strains belonging to the nine human gut species ^31^ (Fig. S1F). Most of the identified sRNAs are highly conserved in *S. copri* (former *P. copri* clade A, n=186) with over 60% of the sRNAs being present in at least two additional *Segatella* species (Fig. S1F). In the case of SrcF, it is present in 97% of all analyzed *S. copri* strains (Fig. 4F) and this high conservation of SrcF allows a deeper examination of the expression of this sRNA in a larger number of human samples carrying uncharacterized *S. copri* isolates. The expression of *S. copri*-encoded sRNAs was analyzed in the publically available metatranscriptomes from humans carrying *S. copri* ^35^ (>5% of *S. copri* reads, n=93, see methods) (Fig. 4G). The median expression of SrcF in those human samples was among the highest expressed sRNAs (1000 TPM, Fig. 4G) (Supplementary Table S2E). Thus, SrcF is a strongly conserved sRNA among *S. copri* strains and is highly expressed in human microbiomes suggesting an important role for this sRNA in the gut.

### SrcF is essential for *S. copri* gut colonization

The structure prediction of the 272 nt SrcF sRNA shows a highly structured 5’end and stem-loop at the 3’end resembling features of canonical non-coding regulatory RNAs in bacteria (Fig. 5A). To test whether SrcF is an important non-coding RNA regulator of gut colonization, we genetically deleted SrcF in the HDD04 strain (Δ*SrcF*) and validated the lack of expression by northern blot detection (Fig. 5B). *S. copri* wild-type (Wt) and *S. copri* Δ*SrcF* strains display a similar growth profile ruling out a growth defect due to the deletion of SrcF *in vitro* (Fig 5C). Next, we colonized OMM12 mice with HDD04 Wt and Δ*SrcF* (ratio 1:1), and monitored the abundance of *S. copri* over time in fecal samples (n=6) (Fig. 5D). Stable colonization of the OMM12 mice by *S. copri* HDD04 (Wt + Δ*SrcF*) was validated via 16S rRNA amplicon sequencing from feces (Fig. 5D) (Supplementary Table S3B). The relative abundance of *S. copri* Wt and Δ*SrcF* was determined by qPCR with strain-specific primers at day 3, 5, 7, and 14 post-colonization revealing that remarkably *S. copri* Wt outcompeted Δ*SrcF* by >1000-fold in abundance by day 14 (Fig. 5E). Consistent with the high expression of SrcF in human microbiomes (Fig. 4), SrcF is an essential factor for *S. copri* gut colonization.

**Figure 5.**
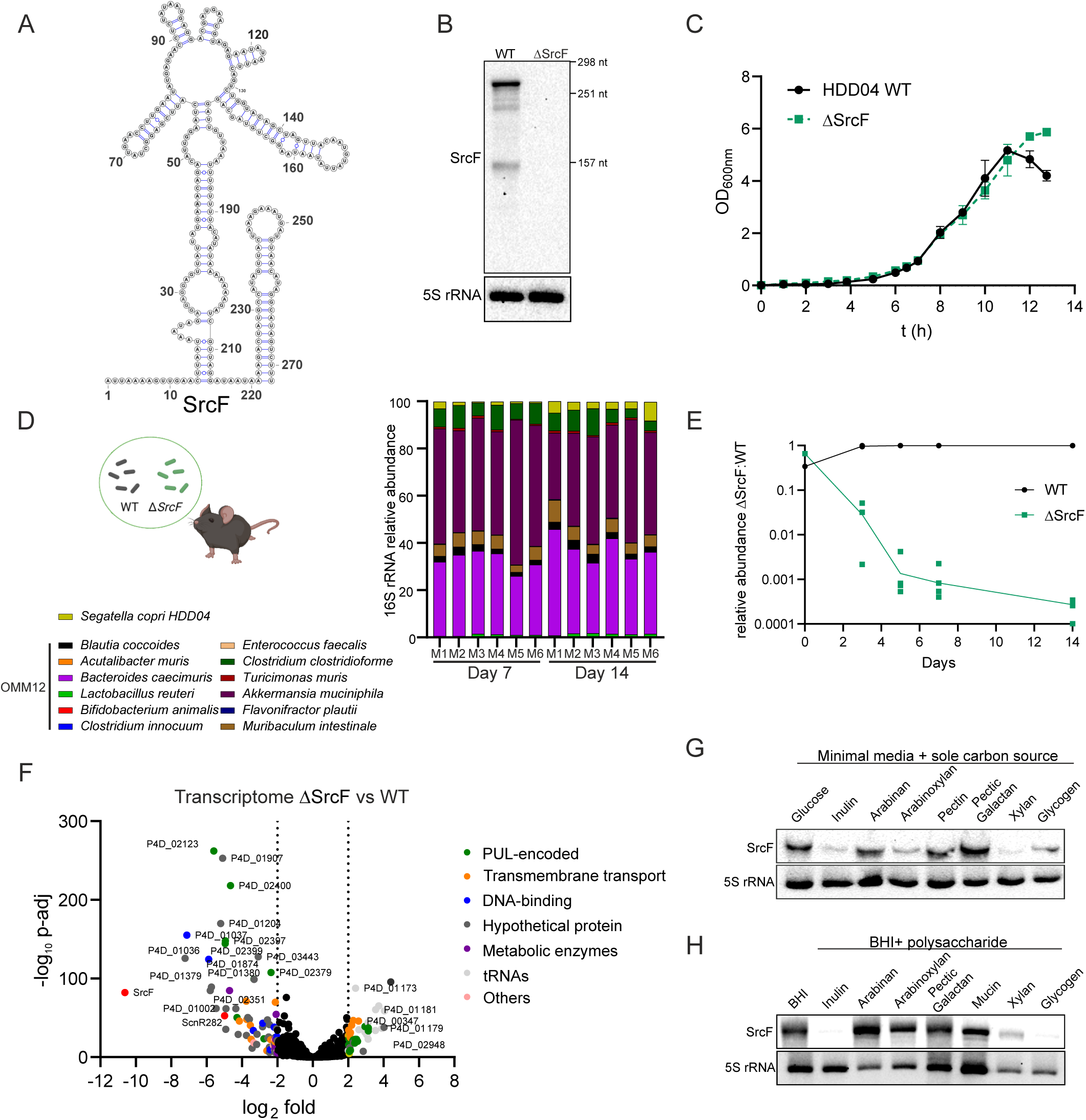
SrcF is essential for *S. copri* gut colonization. Predicted secondary structure of SrcF sRNA. **B.** Northern blot detection of SrcF in *S. copri* Wt and *S. copri* ΔSrcF grown to stationary phase. 5S rRNA was detected as a loading control. **C.** Growth curve of *S. copri* Wt and *S. copri* ΔSrcF in BHI media. **D.** Relative abundance of *S. copri* (Wt+ ΔSrcF) and the Oligo-Mouse-Microbiota (OMM12) members. 16S rRNA amplicon sequencing was carried out from genomic DNA extracted from fecal samples at day 7 and day 14 post *S. copri* gavage. **E.** Relative abundance of *S. copri* Wt and *S. copri* ΔSrcF in fecal samples of OMM12 mice (n=6) colonized with *S. copri* Wt and ΔSrcF (1:1). Samples were collected at day 0 (input), day 3, day 5, day 7, and day 14. **F.** Volcano plot representing the log2 fold-change of gene expression in *S. copri* ΔSrcF compared to *S. copri* Wt grown to stationary phase in BHI *in vitro*. The putative gene function of differentially regulated genes is indicated. **G.** Northern blot detection of SrcF in *S. copri* Wt grown in minimal media in the presence of the indicated sole carbon source. Samples were collected at OD_600nm_ 0.5. 5S rRNA was detected as a loading control. **H.** Northern blot detection of SrcF in *S. copri* Wt grown in BHI supplemented with the indicated carbon source. Samples were collected upon 20 h incubation. 5S rRNA was detected as a loading control.

### SrcF orchestrates metabolic rewiring in response to specific complex polysaccharides

The deletion of SrcF leads to a prominent fitness defect in *S. copri* gut colonization presumably via deregulating gene expression. To gain an insight into the genetic network under the regulation of SrcF, we carried out a gene expression analysis of *S. copri* Wt and Δ*SrcF* grown to Stat when SrcF expression is the highest (see Fig. 2C) (Fig. 5F). SrcF deletion led to the upregulation of 42 genes, and the downregulation of 69 genes (-2>log_2_ fold >2, *p*-adj <0.05) (Fig. 5F) (Supplementary Table S3D). In the absence of SrcF, PUL11, PUL22, and PUL23 in *S. copri* HDD04 are upregulated, while PUL16 is downregulated suggesting a role for SrcF in regulating PUL expression (Fig. 5F). In addition, the expression of genes encoding for putative transport systems (e.g P4D_01551-1555, encoding for outer membrane and inner membrane transporters) and metabolic enzymes (e.g P4D_02317-2319, encoding for beta-galactosidase and alpha_L-fucosidase) are also deregulated in Δ*SrcF* suggesting that it could be involved in the regulation of substrate uptake and metabolism (Fig. 5F, Supplementary Table S3D). DNA-binding proteins are also downregulated in the absence of SrcF as well as a subset of genes of unknown function (Fig. 5F). Altogether, the absence of SrcF resulted in substantial deregulation of the *S. copri* transcriptome, particularly, of pathways involved in the regulation of the metabolism and nutrient acquisition. This suggests that SrcF could be mediating metabolic rewiring of *S. copri* at the gene expression level.

As SrcF deletion altered the expression of multiple PULs and nutrient acquisition genes, we hypothesized that the expression of SrcF itself could be influenced by the availability of carbon sources. *S. copri* was grown in minimal media with complex carbohydrates known to be utilized as sole carbon sources by *S. copri* ^21^, namely inulin, arabinan, arabinoxylan, pectin, pectic galactan, xylan, and glycogen; Glucose served as a simple sugar control and expression of SrcF was determined. Notably, SrcF is poorly expressed when *S. copri* is grown in inulin, xylan, arabinoxylan, and glycogen as sole carbon sources (Fig. 5G). Similarly, the supplementation of inulin, xylan, and glycogen to rich media also led to the downregulation of SrcF expression (Fig. 5H). Overall, SrcF levels are modulated by the metabolization of complex carbohydrates utilized by *S. copri*, which in turn would impact metabolic gene expression under SrcF control.

### SrcF is repressed by inulin metabolization

To better understand the specific regulation of SrcF expression by complex polysaccharide availability, we explored how inulin utilization by *S. copri* affects SrcF expression. *S. copri* HDD04 encodes for 29 predicted PULs, of which only 4 have been experimentally validated for the utilization of a specific complex polysaccharide ^21^. It remains unknown which PULs are required for the utilization of xylan and glycogen, however, we have previously identified PUL26 as essential for the utilization of inulin ^21^. PUL26 expression is controlled by a hybrid two-component system (HTCS) regulator (*HTCS^Inu^*) that is required for *S. copri* growth on inulin as a sole carbon source ^21^ (Fig. S2A).

We asked whether the inulin β-(2-1)-fructose polymer or the derived oligosaccharides and monosaccharides lead to the downregulation of SrcF. For this purpose, *S. copri* was grown in BHI supplemented with inulin, fructooligosaccharides (FOS), or fructose, and the expression of SrcF was determined by northern blot (Fig. 6A). Interestingly, all three inulin components including the monosaccharide fructose lead to the downregulation of SrcF (Fig. 6A). Comparing the expression of SrcF between Wt and Δ*htcs^Inu^* showed that the downregulation of the sRNA by inulin and FOS was lost in the Δ*htcs^Inu^* background (Fig. 6B). Interestingly, providing the inulin-derived monosaccharide fructose directly also failed to repress SrcF in Δ*htcs^Inu^* suggesting that fructose-mediated SrcF repression is signaled through HTCS^Inu^ (Fig. 6B). In agreement, the addition of fructose leads to the repression of SrcF in the Wt but not the Δ*htcs^Inu^* strain. However, the sRNA is not repressed in either strain in the presence of other monosaccharides such as glucose, galactose, and xylose (Fig. S2B). Of note, arabinose repressed SrcF and might represent an additional signal that impacts SrcF expression (Fig. S2B). The HCTS regulator in PULs dedicated to fructan (inulin or levan) degradation in the related *Bacteroides spp.* binds directly to fructose to activate PUL expression ^36^. Interestingly, HTCS in *S. copri* PUL26^Inu^ displays a high conservation with HTCS^Inu^ from *Bacteroides* including identified residues that interact with fructose ^36^ suggesting that the *S. copri* HTCS^Inu^ might interact directly with fructose derived from inulin degradation in *S. copri* (Fig. S3).

**Figure 6.**
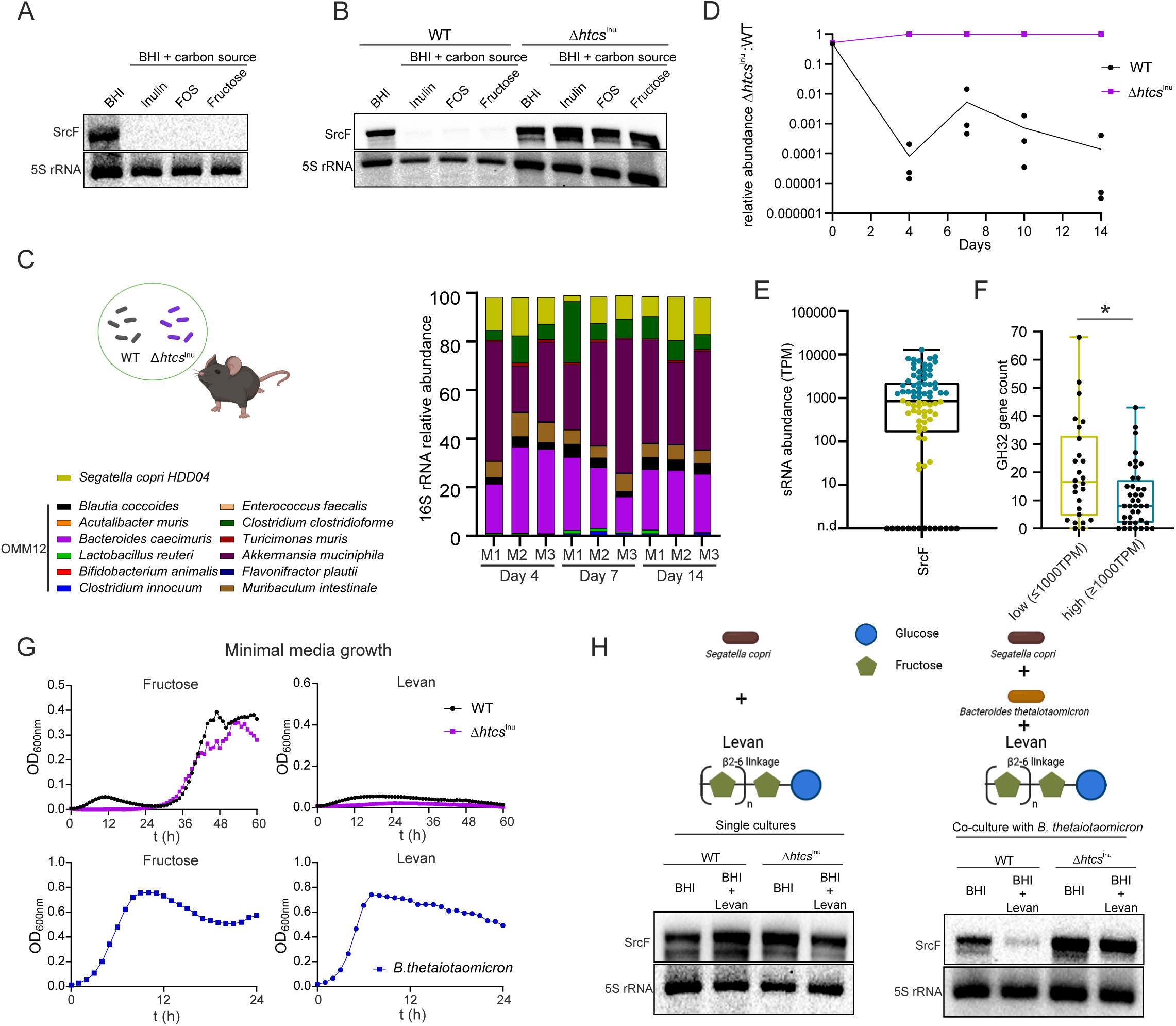
SrcF is repressed by inulin metabolization by *S. copri* and by polysaccharide degradation by cohabiting commensals. Northern blot detection of SrcF in *S. copri* Wt grown in BHI-S supplemented with inulin, fructooligosaccharides (FOS) or fructose. Samples were collected upon 20 h incubation. 5S rRNA was detected as a loading control. **B.** Northern blot detection of SrcF in *S. copri* Wt and *S. copri* Δ*htcs^Inu^* grown as in A. **C.** Relative abundance of *S. copri* (Wt+ Δ*htcs^Inu^*) and the Oligo-Mouse-Microbiota (OMM12) members. 16S rRNA amplicon sequencing was carried out from genomic DNA extracted from fecal samples on day 4, day 7, and day 14 post *S. copri* gavage. **D.** Relative abundance of *S. copri* Wt and *S. copri* Δ*htcs^Inu^* in fecal samples of OMM12 mice (n=3) colonized with *S. copri* Wt and Δ*htcs^Inu^* (1:1). Samples were collected at day 0 (input) day 4, day 7, day 10 and day 14. **E.** Expression of SrcF in human microbiomes carrying *S. copri* (n=82). The median of TPM expression is represented. High SrcF-expressing samples (>1000 TPM) are indicated in blue and low SrcF-expressing samples (<1000 TPM) are indicated in yellow. **F.** Abundance of glycosyl hydrolase 32 (GH32) in high and low SrcF-expressing microbiomes. Wilcoxon, p = 0.019. **G.** Growth curve of *S. copri* Wt, *S. copri* Δ*htcs^Inu^* and *Bacteroides thetaiotaomicron* VPI-5482 in minimal media supplemented with fructose or levan as sole carbon source. **H.** Northern blot detection of SrcF in *S. copri* Wt and *S. copri* Δ*htcs^Inu^* grown in BHI-S, and in BHI-S supplemented with levan in single cultures (upper panel) or in co-culture with *B. thetaiotaomicron* (bottom panel). Samples were collected upon 20 h incubation. 5S rRNA was detected as a loading control.

Because SrcF is essential for colonization and its expression can be repressed by PUL26 HTCS^Inu^, we hypothesized that PUL26 HTCS^Inu^ could impact *S. copri* gut colonization. *S. copri* Wt and Δ*htcs^Inu^* were competed (1:1 ratio) in OMM12 mice and colonization was verified by 16S rRNA amplicon sequencing from feces (Fig. 6C) (Supplementary Table S3C). Strikingly, *S. copri* Δ*htcs^Inu^* outcompeted the Wt strain indicating that inulin utilization is not essential for *S. copri* colonization (Fig. 6D). On the contrary, deletion of Δ*htcs^Inu^* confers a colonization advantage in the gut as Δ*htcs^Inu^* strongly outcompetes the Wt strain (>1000-fold) (Fig. 6D). This colonization advantage of Δ*htcs^Inu^* could be explained by the insensitivity of this strain to the fructose-mediated SrcF repression (Fig 6D). Overall, these results show that fructan utilization by *S. copri* downregulates the essential colonization factor SrcF, and abolishing inulin utilization provides a colonization advantage to *S. copri* in the gut of gnotobiotic mice. In agreement, transcriptomic analysis of *S. copri* in the mouse gut indicates that PUL26 is downregulated in the colon and the cecum of OMM12-*S. copri* colonized mice while other PULs such as PUL14, PUL21, and PUL24 that are required for arabinan, pectic galactan, and arabinoxylan utilization ^21^ are induced in the mouse gut compared to when grown *in vitro* (Fig. S4) (Supplementary Table S3E-F).

### Polysaccharide degradation by cohabiting commensals impacts *S. copri* gut colonization factor expression

SrcF expression is repressed by inulin utilization, and it is largely highly expressed in human gut microbiomes carrying *S. copri*. We asked whether polysaccharide degradation capacity in human microbiomes, particularly that of fructans, would recapitulate the observed regulation *in vitro*. SrcF is not homogeneously expressed in the examined human stool, with 40 samples (43%) displaying higher SrcF expression (>1,000 TPM, high) and, 28 samples (30%) displaying lower SrcF expression (<1000 TPM, low) (Fig. 6E). A subset of 25 samples (27%) displayed no SrcF expression (TPM=0) potentially due to either insufficient coverage or absence of this locus in those samples (Supplementary Table S2E). SrcF expression in human donors does not correlate with *S. copri* abundance (R=-0.018, p=0.88; Fig. S5A), suggesting that extrinsic signals such as microbiota differences may contribute to the SrcF expression profile. Taxonomic analysis of the microbiome composition displays no significant differences in alpha-diversity between high and low SrcF expression samples while minor differences in beta-diversity are observed (p=0.011) (Fig. S5B-C). Functional analysis based on the presence of carbohydrate-active enzymes (CAZymes) suggests a distinct functionality in high and low SrcF-expressing microbiomes (p=0.001; Fig. S5D, Supplementary Table S4). Notably, the gene content of GH32, the glycosyl hydrolases responsible for fructan cleavage, was significantly lower in high SrcF samples compared to low SrcF samples (Fig. 6F). This observation led us to hypothesize that fructan metabolization by other commensals in the communities could contribute to SrcF expression and consequently affect *S. copri* colonization. Of note, distinct fructan utilization has been shown to drive competition and cooperation among *Bacteroides* species, where *Bacteroides caccae*, *Bacteroides ovatus,* and *Bacteroides fragilis* utilize inulin (β2-1 fructan), but not levan (β2-6 fructan), whereas *B. thetaiotaomicron* in turn utilizes levan but not inulin ^36^. In the case of *S. copri,* while it can utilize inulin, it cannot grow on levan as the sole carbon source (Fig. 6G), and therefore levan presence should not affect SrcF expression. In agreement, levan supplementation failed to downregulate SrcF expression in *S. copri* Wt or Δ*htcs^Inu^* (Fig. 6H). When *S. copri* Wt was co-cultured with *B. thetaiotaomicron*, SrcF was downregulated in a levan-dependent manner demonstrating that degradation of levan by *B. thetaiotaomicron* leads to downregulation of SrcF in *S. copri* (Fig. 6H). Consistently, the presence of *B. thetaiotaomicron* and levan did not lead to SrcF downregulation in Δ*htcs^Inu^*, supporting the model in which fructose-mediated downregulation of SrcF is signaled through HTCS^Inu^ (Fig. 6H, S6). Altogether, we show that complex polysaccharide degradation might contribute not only to cross-feeding mechanisms ^37^ but also to impact gene expression in cohabitating commensals.

## DISCUSSION

Bacterial fitness depends on the ability to sense and rapidly respond to changing environments to survive. In the gut, commensal bacteria must effectively interpret and respond to signals from the host, from nutrients in the diet, as well as engage in both cooperative and antagonistic interactions with other inhabitants of the gut to achieve successful and stable colonization. While understanding these regulatory mechanisms is of great importance, both for fundamental research and the development of microbiome-based therapeutics, studies of commensal gene regulation involved in intricate colonization strategies for many prevalent non-model human commensals have been scarce.

*S. copri* is one such prevalent member of the human gut microbiome that is estimated to have colonized predominantly humans for hundreds of thousands of years ^5^. The recently expanded strain collections of members of the *S. copri* complex and the development of genetic tools have enabled the initiation of functional studies of *S. copri* ^21,31^. In this study, we uncovered a largely similar global RNA architecture for both *S. copri* HDD04 and DSM18205^T^, which revealed a wealth of non-coding sRNA regulators conserved in *Segatella copri* (former *Prevotella copri* clade A) (Fig. 2, S1). Due to the observed strain-to-strain differences in *S. copri* genetic tractability have led us to propose the *S. copri* HDD04 as a model for future functional studies as it is substantially more genetically tractable than the type strain *S. copri* DSM18205^T 21^. The high conservation of the identified sRNAs and the general high abundance of *S. copri* in individuals that carry the species allows the expression analysis of these regulatory RNAs directly in human microbiomes. Using this approach, we identified an essential RNA colonization factor, ScrF, which is generally highly expressed in human microbiomes.

Deletion of SrcF strongly impacts the ability of *S. copri* to colonize the gut and deregulates an important number of genes putatively involved in both the uptake and utilization of nutrients, including PULs (Fig. 5). *S. copri* is a gut bacterium that is well-equipped to metabolize complex carbohydrates ^5,10,20^. The competition for varying carbon sources in the gut requires versatile fine-tuning of gene expression for efficient energy harvesting and successful colonization ^2,22^. For model organisms such as *E. coli*, we have a deep understanding of energy source hierarchies, and sRNAs have been shown to play a role in reshaping bacterial gene expression in response to carbon source availabilities ^38,39^. However, which carbon sources are preferred and how hierarchies are established at the transcriptomic level remain unknown for most commensal bacteria, including *S. copri*. In addition to being a regulator of PUL gene expression, SrcF levels are also modulated by the presence of complex carbohydrates suggesting that SrcF might represent a regulatory hub that integrates nutrient cues essential for *S. copri* survival in the gut. PUL expression is tightly controlled and is frequently regulated by hybrid two-component systems (HTCS) that sense oligosaccharides of two to eight subunits resulting from polysaccharide breakdown ^2,40,41^. In *B. thetaiotaomicron,* the HTCS regulator of the PUL that breaks down the fructan levan has been shown to sense directly fructose monosaccharides ^36^. In *S. copri,* we have identified that fructose downregulates SrcF expression and that the signaling is dependent on the HTCS^Inu^ regulator of the PUL26. In agreement, conservation analysis suggests HTCS^Inu^ could bind directly to fructose in *S. copri* (Fig. S3). How exactly HTCS^Inu^ regulates ScrF expression remains to be elucidated.

Fructans are common plant-derived polysaccharide in our diet and their metabolism plays a pivotal role in the competition and cooperation among Bacteroidota ^36,37^. It drives competition between *B. caccae* and *B. thetaiotaomicron* in mice ^36^, and surface breakdown of inulin by *B. ovatus* leads to cross-feeding with *Phocaeicola vulgatus*^37^. Similarly, the extracellular breakdown of levan by *B. thetaiotaomicron* supports the growth of other *Bacteroides spp* ^42^. We showed that the degradation of levan by *B. thetaiotaomicron* leads to SrcF repression in *S. copri* (Fig. 6). Concurrently, we found that the fructan-specific glycosyl hydrolase GH32 correlates with lower expression of SrcF in human microbiomes, which suggests a higher capacity of these microbiota to breakdown fructans and release fructose. Altogether, the breakdown of complex polysaccharides not only contributes as an energy resource for gut commensal bacteria but could serve as an important inter-bacterial signal that impacts gene expression and colonization competence. Interestingly, monomeric fructose and glucose from the diet have been shown to affect the expression of Roc, a regulator of colonization in *B. thetaiotaomicron* ^43,44^. Notably, the 5’mRNA leader of *roc* is required for the response to fructose and glucose suggesting a post-transcriptional mediated control of colonization in *B. thetaiotaomicron*^43^. While SrcF is not present in *B. thetaiotaomicron*, this example highlights the potential of post-transcriptional regulation in the control of colonization of commensals in the gut, particularly in response to dietary components.

Most sRNAs impact gene expression via interaction with mRNA targets aided by RNA binding proteins (RBPs) that can lead to an upregulation or downregulation of the target gene ^45^. Deletion of SrcF sRNA impacts strongly *S. copri* gene expression, whether SrcF functions in conjunction with an RBP and whether it regulates directly or indirectly target genes remains to be determined. Based on homology, *S. copri* lacks all known small RNA binding proteins. Specifically, it lacks the three main RBPs described Hfq, ProQ, and CsrA in bacteria ^46^ as well as the recently described new RBP family in *B. thetaiotaomicron* ^47,48^. Of note, genetics of *S. copri* continue to be challenging, and to date, we lack genetic systems that allow overexpression or ectopic expression of genetic constructs from a neutral genomic site as the pNBU system in *Bacteroides spp*, which are instrumental for genetic dissection ^49–54^. Nevertheless, the identification of TSSs and regulatory elements in *S. copri* serves as a pivotal stepping stone toward the development of genetic tools, including constructs for ectopic gene expression. Such advancements hold promise for facilitating the overexpression of genes ^50,52^, including sRNAs, or adapting CRISPRi-based screens ^55–57^ to streamline gene function analyses.

To functionally study *S. copri* in the gut, we employed the reduced mouse community OMM12, stable across multiple generations and consistently generated in various animal facilities ^34,58^. Colonizing OMM12 mice with *S. copri* allowed us, for the first time, to conduct dRNA-seq of commensal bacteria in the gut and identify TSSs of genes expressed *in vivo* in a single experiment, eliminating the need to expose the bacteria to multiple stimuli *in vitro* to mimic *in vivo* conditions. While *S. copri* exposure to different dietary components and/or microbiota communities of higher complexity might uncover additional elements of the *S. copri* RNA landscape, our current approach allowed the identification and validation of the first colonization factor in *S. copri.* Future studies aiming at the establishment of genome-wide screens in *S. copri* should warrant further discovery of gut colonization requirements of *S. copri*.

Finally, microbiome composition plays a prominent role in human health and a long-standing goal has been to identify and modify specifically problematic microbiota members. Here, we identified an *S. copri*-specific intergenic non-coding sRNA that is essential for gut colonization. *S. copri* presence has been associated with the onset of rheumatoid arthritis in humans ^12,15^ and certain *S. copri* isolates have been shown to exacerbate the disease in a mouse model ^16^. SrcF presents an ideal gene target for the precision editing of microbiomes carrying problematic *S. copri* strains. This could constitute a proof-of-principal for other human gut commensals and identify additional promising non-coding RNA targets for microbiome intervention. Moreover, the presence of certain *Prevotella copri* strains (now *S. copri*) is a principal contributor to the success of microbiome-directed complementary food treatment to fight malnutrition in Bangladeshi children ^59,60^. Indicating that understanding the factors that drive successful *Segatella* spp colonization could have a tremendous impact on future and current microbiome intervention therapies.

## MATERIALS AND METHODS

### Bacterial growth

*Segatella copri* strains and derivative strains (Supplementary Table S5) were grown in BHI-S liquid media composed of Brain Heart infusion (BHI) broth (Oxoid, #CM1135B) (37g/l) supplemented with 10% FBS (Sigma Aldrich, #F7524) and vitamin K3 at 1 μg/ml. For solid growth, strains were grown on BHI agar plates (37 g/l BHI, 18 g/l agar) supplemented with 5 % of horse blood (Thermo Scientific, #SR0050C) and vitamin K3 at 1 μg/ml. Incubation was carried out at 37 °C in an anaerobic chamber (Coy Laboratory Products) with an atmosphere of 10% CO_2_, 4% H_2_, and 86% N_2_. For liquid growth, *S. copri* was inoculated from colony and grown overnight in BHI-S standing cultures at 37°C, the overnight culture was subcultured 1:100 and *S. copri* was grown to the desired OD_600nm_. When necessary, BHI-S was supplemented with 5 μg/ml erythromycin. *Escherichia coli* strains were cultivated in Lysogeny broth (LB) (tryptone 10 g/l, yeast extract 5 g/l, and sodium chloride 10 g/l). Bacterial cultures were aerobically incubated at 37°C with 200 rpm shaking. When required LB was supplemented with ampicillin (200 μg/ml) and 0.3 mM 2, 6-diaminopimelic acid (DAP).

*Segatella copri* was also grown in the presence of complex polysaccharides as the sole carbon source. It was grown in minimal media prepared as previously described ^21^ supplemented 1:1 with complex polysaccharides solutions at 10 g/l. Alternatively, polysaccharide solutions were supplemented 1:1 to BHI-S. Solutions of arabinan (Megazyme), arabinoxylan (Megazyme), pectic galactan (Megazyme), xylan (carbosynth), mucin (Sigma), glycogen (Sigma Aldrich), pectin (Fluka), levan (Sigma aldrich), and inulin (Carbosynth) were prepared in water and sterilized by autoclaving. Alternatively, when possible, solutions were dissolved in water and sterilized by filtration. BHI-S was also supplemented with oligosaccharides and monosaccharides solutions 1:1. Fructooligosaccharides (Carbosynth), fructose (Sigma), glucose (Sigma), galactose (sigma), xylose (Sigma) and arabinose (Sigma-Aldrich) solutions (10 g/l) were prepared in water and sterilized by filtration.

### Genetic manipulation

Oligonucleotides and plasmids used for genetic manipulation are listed in Supplementary Table S5. *Segatella copri* deletion was generated as previously described ^21^ with minor changes. Shortly, 2 kb fragments upstream and downstream of SrcF were PCR amplified with Q5 High Fidelity DNA Polymerase (New England Biolabs) and cloned in PCR amplified pEx-deletion-ermG via DNA Gibson assembly (HiFi DNA Assembly Master Mix, New England biolabs). Clones were genotyped by PCR and positive clones were validated by Sanger sequencing at Microsynth Seqlab (Germany). Conjugation and selection of gene deletion of SrcF in *S. copri* HDD04 was carried out as previously described. Shortly, *E. coli* β2155 carrying pEx-deletion-ermG-SrcF was grown in LB supplemented with ampicillin in aerobic conditions to OD_600nm_ 0.5-0.7, 1ml of cells was pelleted, washed in LB without antibiotic, and pelleted cells transferred to the anaerobic chamber (Coy lab). *S. copri* HDD04 recipient strain was grown in BHI-S to OD_600nm_ 0.5-0.7. A pellet of the donor strain was resuspended in 100 μl of *S. copri* culture, spotted on BHI-blood agar plates containing DAP (*E. coli* β2155 auxotroph for DAP), and incubated at 37 °C for 18h for conjugation under anaerobiosis. Bacterial growth was resuspended in 1ml of BHI-S and 100 μl plated in BHI-blood agar supplemented with 5 μg/ml erythromycin and 200 μg/ml of gentamycin to select for transconjugants after 2-4 days incubation at 37 °C in anaerobiosis. Correct integration was validated by PCR and correct transconjugants were subcultured in BHI-S for allelic exchange and plated on YT (yeast extract 5 g/l and tryptone 10g/l) agar plates supplemented with 5% sucrose as negative selection of plasmid integration maintenance. Clones grown after 2-4 days incubation at 37 °C were streaked on BHI-blood plates with and without erythromycin and antibiotic sensitive strains were genotyped by PCR for gene deletion and validated by Sanger sequencing at Microsynth Seqlab (Germany).

### Total RNA isolation

Strains of interest were grown anaerobically to the desired cell density and the biomass of 4 units of OD_600nm_ was collected 5:1 (v/v) with stop solution (95% Ethanol, 5% phenol) and snap frozen in liquid nitrogen. Bacterial total RNA was extracted by hot phenol method followed by a DNase treatment. For RNA extraction from mice gut content, upon sacrificing the mice cecum and colon content was immediately resuspended in RNA shield solution (#R2001) and stored at -80 _o_C. Total RNA was extracted with ZymoBIOMICS RNA Mini Kit (#R2001) following manufacturers instruction. Quality of the RNA was determined by bioanalyzer at genome analytics core facility of the Helmholtz Centre for Infection Research (HZI).

### Northern blot and DIG riboprobes

Direct detection of sRNAs was carried out by northern blot. Shortly, 10 μg of total RNA was subjected to electrophoretic separation in Tris-Borate-EDTA (TBE) 6% acrylamide gels containing 8.3 M urea. (300V, 30mA). RNAs were transferred to nylon membrane (Sigma Aldrich, #15356-1EA) and crosslinked to the membrane by UV crosslinking. For probes, full-length anti-sense DIG-labeled RNA probes (riboprobe) complementary to the target sRNA were used. For riboprobe generation, PCR template was obtained for each probe and RNA probes were generated with MAXIscript T7 *in vitro* transcription kit (Thermo Scientific, AM1312) following manufacturers instruction with minor modifications, and DIG RNA labeling mix (Sigma Aldrich, #11277073910) was added as nucleotide mix. Labeled riboprobe were further purified via Microspin G-50 columns (Cytiva, # 27-5330-01). Membranes were hybridized with riboprobes in Roti-HYBRI quick buffer (CarlRoth, #A981.1) at 68 _o_C and washed with 2x SSC and 0.5xSSC. Signal detection was carried with DIG luminescence detection kit (Sigma Aldrich, #11363514910) following manufacturers instruction and signal captured with ChemiDoc (Biorad). For 5S rRNA loading control detection, a 5’ DIG labeled oligonucleotide was used as probe (Eurofins). The membrane was incubated with the 5S rRNA probe at 42 °C and washes carried out with 5xSSC 1xSSC and 0.5x SSC. The signal was detected as for riboprobes. Oligonucleotides used to generate PCR templates for riboprobes are listed in Supplementary Table S5.

### RNA processing and cDNA library preparation for sequencing

For differential RNA sequencing (dRNA-seq), libraries were prepared at Vertis Biotechnologie AG as previously described ^61^. First, the total RNA samples were fragmented using ultrasound (4 pulses of 30 sec at 4°C) followed by a treatment with T4 Polynucleotide Kinase (New England Biolabs). The RNA samples were then split into two halves and one half was subjected to Terminator Exonuclease treatment (+TEX), the other half was left untreated (-TEX). The RNA samples were poly (A)-tailed using poly(A) polymerase. Then, the 5’PPP structures were removed using RNA 5’ polyphosphatase (Epicentre). Afterwards, an RNA adapter was ligated to the 5’-monophosphate of the RNA. First-strand cDNA synthesis was performed using an oligo(dT)-adapter primer and the M-MLV reverse transcriptase. The resulting cDNAs were PCR-amplified to about 10-20ng/µl using a high-fidelity DNA polymerase and purified using the Agencourt AMPure XP kit (Beckman Coulter Genomics). Pooled libraries were sequenced on an Illumina NextSeq 500 platform at the Core Unit SysMed at the University of Würzburg for bacterial *in vitro* samples and on an Illumina Novaseq at the genome analytics core facility of the Helmholtz Centre for Infection Research (HZI) for mouse derived samples.

For conventional RNA sequencing, total RNA from *S. copri* strains was subjected to bacterial ribosomal RNA (rRNA) depletion by the use of Pan-prokaryote riboPOOL Kit (siTOOLs Biotech) following manufacturer’s instructions. Libraries for Illumina sequencing were prepared using the NEBNext Ultra Directional RNA Library Prep Kit for Illumina (New England Biolabs) following manufacturer instructions. For metatranscriptomic sequencing, total RNA from mouse gut or human donors was subjected to ribosomal RNA depletion by Ribo-Zero Plus Microbiome rRNA Depletion Kit (Illumina) and cDNA libraries were generated by using Illumina Stranded Total RNA Prep (Illumina) following manufacturer’s instructions.

### Differential RNA sequencing (dRNA-seq) data analysis

Libraries for differential RNA sequencing (+/- TEX) were sequenced with a depth of 15 million reads/library. Samples from early exponential phase (EEP), mid exponential phase (MEP and stationary phase (Stat) were integrated in the analysis of differential RNA-seq of the respective *S. copri* HDD04 and DSM18205^T^ strains. Generated FASTQ files were mapped to *S. copri* HDD04 or *S. copri* DSM18205^T^ reference genome by using the READemption (1.05) pipeline ^62^ and coverage files were integrated in the ANNOgesic pipeline ^25^ as previously described with all parameters at the default setting ^26,61^. ANNOgesic allows the identification of transcription start sites (TSSs), terminators, transcript length, untranslated regions, small RNAs and promoter motifs. Shortly, TSSs were identified by the use of TSSpredator ^63^ that categorized TSSs in five categories based on its location relative to the annotated CDS. First, primary TSSs are defined as the TSS with the highest coverage within the 300 nt window upstream of a given CDS, the remaining TSSs are defined as secondary. Secondly, internal TSSs are located within a given CDS, while antisense TSSs are located within a CDS or a 100 nt window flanking the CDS in an antisense orientation. Finally, predicted TSSs that do not associate with any CDS, were classified as orphan and might be indicative of intergenic unannotated genetic elements. Combination of TranstermHP ^64^ and RNAfold ^65^ allows ANNOgesic pipeline to predict terminators and therefore boundaries of transcripts ^25^, fragmented library of the respective strains was included in the analysis for terminators predictions and default parameters were used. For promoter putative motif identification, the 50 nt upstream of all identified primary TSSs were extracted and used as input in MEME ^27^ to identify conserved motifs.

For TSS mapping from *S. copri* colonized OMM12 mice, generated libraries were sequenced with 400 million reads sequencing depth paired-end 2x50bp. To obtain *S. copri* mapped reads, reads were aligned to *S. copri* HDD04 genome and to the 12 members of Oligo-Mouse-Microbiota (OMM12) members ^66,67^ by the use of READemption (2.0.0) and --*crossalign_cleaning* command to remove reads mapped to multiple species ^62^. Generated coverage file for *S. copri* only reads were integrated in the ANNOgesic pipeline and TSSpredator to map TSSs with default settings. Coverage file of (+/- TEX) libraries were loaded into Integrative Genomics Viewer ^68^ (IGV) to visualize transcription start sites (TSSs).

### Total RNA sequencing data analysis

Pooled libraries were paired-end (2x50bp) sequenced on an Illumina Novaseq at the genome analytics core facility of the Helmholtz Centre for Infection Research (HZI). Generated FASTQ files were mapped to *S. copri* HDD04 reference genome by using the READemption pipeline ^62^. Gene-wise read counts were normalized to TPM (transcripts per kilobase million) that takes into account sequencing depth (number of mapped reads) and transcripts length. Differentially expressed genes between *S. copri* Wt and *S. copri* ΔSrcF libraries were calculated with DESeq2 ^69^ that utilizes Wald test to determine the P-value and the Benjamini-Hochberg to correct p-values for multiple testing (P-adj) ^69^.

### Metatranscriptomic data analysis

Publicly available metatranscriptomic data from human fecal microbial communities from a cohort of adult men (372 metatranscriptomes) ^35^ was reanalyzed to determine the abundance of newly identified small RNAs in *S. copri*. Raw reads were trimmed for low quality and filtered against the phix174 and human hg19 genome with bbduk. Next, ribosomal reads were removed using ribodetector v0.3 ^70^. To avoid false-positive mapping form close related species, we perform a taxonomic species profiling: all libraries were mapped against the genomes the Unified Human Gastrointestinal Genome collection version 2 ^71^ (n = 4,744) with bwa-mem2. Reads mapped to Clade-A representative genome “*Prevotella* sp900557255” were filtered via samtools into a new read library. 93 of those metatranscriptomes mapped over 5% of the reads to *S. copri* from the Unified Human Gastrointestinal Genome (UHGG) collection ^71^. Filtered reads to *S. copri* were remapped to our *S. copri* HDD04 via using READemption pipeline ^62^ and relative expression of newly annotated intergenic sRNAs determined normalizing reads to (transcripts per kilobase million) TPM.

For human healthy donors of the Mikrodivers, donors cohort a respective *S. copri* strain has been previously isolated from donors A, B, C and D: *S. copri* HDA03, *S. copri* HDB01, *S. copri* HDC01 and *S. copri* HDD04 ^21^. Annotation of *S. copri* genomes were amended with coordinates of conserved newly annotated sRNAs in *S. copri* HDA03, *S. copri* HDB01 and *S. copri* HDC01 (Fig. 2). The reads were filtered as described previously ^21^ and then mapped to the genome annotations of respective *S. copri* isolates using the READemption pipeline ^62^. The relative expression levels of newly annotated intergenic sRNAs were determined by normalizing the read coverage to transcripts per million (TPM). The fold change in sRNA expression among human donors was determined with their respective *S. copri* isolate grown *in vitro* in BHI as reference.

### Genomic DNA extraction and amplicon 16S rRNA sequencing

Genomic DNA from mice feces was extracted by using ZymoBIOMICS^TM^ DNA Miniprep kit following manufacturer’s instructions. For 16S rRNA amplicon sequencing, the V4 region was amplified as previously described ^72,73^. Pooled libraries were sequenced on an Illumina MiSeq platform, paired-end 250 bp sequencing. Sequenced libraries were processed using QIIME v1.8.0 ^74^. Reads were clustered into 97 % ID OTUs by open-reference OTU picking and representative sequences were determined by use of UPARSE algorithm ^75^ by using the 16S rRNA genes of Oligo-Mouse-Microbiota (OMM12) genomes ^67^ and *S. copri* HDD04.

### Human donors metagenomes library preparation

Genomic DNA from donor A-D stool samples was extracted by using ZymoBIOMICS^TM^ DNA Miniprep kit following manufacturer’s instructions. Genome libraries were prepared using the Illumina DNA PCR-Free Library Kit and the IDT for Illumina DNA/RNA UD Indexes for previously isolated DNA. The libraries were prepared following the manufacturer’s instructions. Quantification of library concentrations was performed using Qubit ssDNA Assay Kit followed by further quantification with KAPA Library Quantification Kit for Illumina. Sequencing was performed on the NovaSeq S4 PE150 platform targeting a depth of 4 Gbp per sample.

### Metagenomes data analysis

The relative abundances of *Segatella* species in the gut microbiome of the four human donors A-D was determined. To estimate relative abundances of species belonging to the *Segatella* genus, a reference-based approach MetaPhlAn 4 (v 4.0.0) ^76^ was applied with default settings to four shotgun metagenomic samples derived from donor A, B, C and D. As former taxonomic labels of *Segatella* species are still in use in the current version of MetaPhlAn 4, we manually updated the labels to their formal taxonomic nomenclature as previously carried out ^4,31^.

For the microbiomes of publicly available data, we examined the association of *S. copri* SrcF expression level with the human gut microbiota of the respective donor. We downloaded 68 shotgun metagenomic samples (Supplementary Table S6B) for which SrcF expression was detected (i.e., TPM > 0) in the corresponding shotgun metatranscriptomic samples as described above ^77^. We first assessed the relative abundance of *S. copri* in the downloaded samples, aligning metagenomic reads of each sample against Clade-A representative genome “Prevotella sp900557255” using bowtie2 (v2.3.5.1; --end-to-end --no-unal -U -S) ^78^. The resulting BAM files were generated with samtools (v1.3.1; view -Sb) and were further cleaned using a python package CMSeq (https://github.com/SegataLab/cmseq) with following criteria: (1) reads alignment quality ≥ 30, (2) reads coverage depth ≥ 5X, (3) minimum identity of reads ≥ 96%, (4) aligned read length ≥ 90 nt, (5) minimum dominant allele frequency ≥ 50%. The abundance of Clade-A representative genome in each sample was then calculated by dividing the cleaned aligned reads by the total number of reads of a metagenomic sample. This approach has been used widely in other studies for extracting strain-specific reads^5,31,79^.

For quantifying the whole microbial community composition of each sample we used MetaPhlAn 4 as described above. Alpha diversities were calculated using Shannon index estimated by R package mia (https://github.com/microbiome/mia) and beta diversities were calculated using Bray-Curtis dissimilarity based on species-level relative abundances, followed by a principal coordinates analysis (PCoA) for the visualization (Python package scikit-bio v0.5.7; http://scikit-bio.org/). The R package vegan v2.6.2 (https://github.com/vegandevs/vegan) was used to perform permutational multivariate analysis of variance (PERMANOVA) for beta diversities.

Moreover, for each sample we also profiled the CAZymes functionality carriage using a custom pipeline. Briefly, we first downloaded the CAZyme database (version 09242021, https://bcb.unl.edu/dbCAN2/download/CAZyDB.09242021.fa). Next, for each metagenomic sample, paired-end reads were merged in one FASTQ file, and then were aligned to the downloaded CAZyme database using PALADIN with parameters of –C -n -a ^80^. The output from PALADIN was converted to a BAM file using samtools (v1.3.1) with settings of view -Sb ^81^. The coverage of the alignments of reads against CAZymes were estimated using an utility script pileup.sh from BBMap (https://jgi.doe.gov/data-and-tools/software-tools/bbtools/bb-tools-user-guide/bbmap-guide/). A CAZyme was considered present in a metagenomic sample only when ≥ 10X coverage depth and 100% coverage breadth were both reached. Jaccard distances based on the presence and absence of CAZymes were calculated and visualized by a principal coordinates analysis (PCoA) (Python package scikit-bio v0.5.7; http://scikit-bio.org/). Statistical significance of the difference between low- and high-SrcF-expression groups in CAZyme sequence abundance was assessed by the Wilcoxon Rank Sum test corrected for multiple hypothesis testing by Benjamini-Hochberg method.

### Conservation analysis of small RNAs in *Segatella* species

The conservation of 27 sRNAs in *Segatella* species was assessed by leveraging 1,135 previously reported high-quality metagenome-assembled genomes (MAGs) which were classified into nine human *Segatella* species (Supplementary Table S6A) ^31,76^. To this end, we tailored a validated method which was previously used to quantify the prevalence of CAZymes in distinct species using MAGs ^5^. Briefly, blastn (v2.5.0) 5 was used to align 27 sRNAs against each of MAGs with custom settings of -word_size 11 -qcov_hsp_perc 60 - perc_identity 70 and other parameters were kept as default. The conservation of each sRNA in *Segatella* species was quantified as the percentage of MAGs in that species for which at least one blast hit was observed.

### *S. copri* colonization of Oligo-Mouse-Microbiota (OMM12) mice

Oligo-MM12 C57BL/6NTac ^34^ mice were bred in isolators (Getinge) in the germfree facility at the Helmhotz Centre for Infection Research. Female and male mice between 8-16 weeks of age were used. Sterilized food and water was provided *ad libitum*. Mice were kept under strict 12-hour light cycle and housed in groups of up to three mice per cage. All mice were euthanized by asphyxiation with CO_2_ and cervical dislocation.

For *S. copri* colonization, *S. copri* WT was grown anaerobically to OD_600nm_ of 0.5-0.8. The culture was transferred to air-locked vials and used for subsequent colonization. The mice were gavaged orally with 200 μl of *S. copri* and anally administered 50 μl of the culture. For *S. copri* colonization competition assays, *S. copri* WT, *S. copri* Δ*SrcF* and *S. copri* Δ*htcs^Inu^* were grown anaerobically in BHI-S to an OD_600nm_ of 0.5-0.8. Strains were mixed (1:1) and the mix (input) transferred to air-locked vials for subsequent colonization. The mice were gavaged as above.

### Competition assays quantitative PCR

The relative abundance of *S. copri* WT and mutant strains in competition assays was monitored at the indicated time points via quantitative PCR (qPCR) using strain specific primers. Oligonucleotides used are listed in Supplementary Table S5. Genomic DNA was extracted from mouse feces as described above at the indicated time points and quantitative PCR was carried out in a CFX96 instrument (BioRad) using KAPA Sybr Fast Universal qPCR master mix (Kapa biosystems (#KK4602). Mean strain DNA quantities were calculated using a standard curve to determine relative abundance of *S. copri* WT, *S. copri* Δ*SrcF* and *S. copri* Δ*htcs^Inu^* in the respective competition experiments.

### Human data collection

The study involving human participants was reviewed and approved by the Ethics commission of the Hannover Medical School (MikroDivers: 8628_BO_K_2019). All patients/participants provided their written informed consent to participate in this study. The MikroDivers cohort is an ongoing study recruiting healthy volunteers aged 18 and above. The inclusion criteria are the willingness and ability to sign the informed consent, to provide a fecal sample, and to answer a questionnaire on geographic origin, diet preferences, age, gender, BMI, weight, and height. The exclusion criteria were recent antibiotic intake (<4 weeks), intake of chemotherapeutics and immunosuppressive medications (ever), acute and chronic inflammatory bowel disease, colon cancer, recipient of fecal microbiota transplants, and being involved in direct patient care (e.g., in a hospital). Until September 2023 more than 30 individuals were included.

### Ethics statement

All animal experiments have been performed in agreement with the guidelines of the Helmholtz Centre for Infection Research, Braunschweig, Germany, the National Animal Protection Law [Tierschutzgesetz (TierSchG) and Animal Experiment Regulations (Tierschutz-Versuchstierverordnung (TierSchVersV)], and the recommendations of the Federation of European Laboratory Animal Science Association (FELASA). The study was approved by the Lower Saxony State Office for Nature, Environment and Consumer Protection (LAVES), Oldenburg, Lower Saxony, Germany; License 19/3267 PrevDiet.

### Data availability

All sequencing data has been deposited under NCBI-SRA Bioproject: PRJNA1082502.

## Acknowledgments

We thank Vertis Biotechnologie AG for excellent technical assistance in generating libraries for dRNA-seq. We thank Agata Bielecka and Achim Gronow for library preparation for RNA-sequencing and 16s rRNA sequencing respectively. We thank Haiyi Wang for technical assistance. We also thank Eric Galvez for processing of Mikrodivers human donors samples. We thank the Core Unit SysMed at the University of Würzburg and the genome analytics core facility of the Helmholtz Centre for Infection Research (HZI) for RNA-seq data generation. Y-EM was supported by a Walter Benjamin position from the Deutsche Forschungsgemeinschaft (DFG).

