## Supplementary Information for "The RNA landscape of the human commensal *Segatella copri* reveals a small RNA essential for gut colonization"

#### Supplementary material

#### Supplementary Figures legends

**Supplementary Figure S1. *S. copri* sRNA annotation. A.** Northern blot detection of housekeeping non-coding RNAs tmRNA, M1RNA, and 6S RNA in S. copri DSM18205. On the left, the genomic location is indicated. On the right detection of expression from total RNA samples from EEP, MEP, and Stat growth conditions. 5S rRNA was detected as a loading control. **B.** Sequence length of annotated non-coding small RNAs (intergenic and mRNA-derived). In green indicated transcripts that encode for a small ORF of under 50 aminoacids. **C.** Northern blot detection of ScnR157 in *S. copri* HDD04 DSM18205$^T$ and four additional *Segatella spp* grown in BHI to stationary phase. 5S rRNA was detected as a loading control. **D.** Genomic location of ScnR157 in *S. copri* and other *Segatella spp*. **E.** Alignment of the nucleotide sequence for ScnR157 and the upstream 50 nt among *Segatella spp.* strains. High conservation of the sRNA sequence and canonical *Segatella* promoter motif are indicated in blue. **F.** Heat map of intergenic sRNAs conservation in *Segatella* spp. Number of strain genomes investigated is indicated in brackets for each of the nine species of *Segatella* species complex associated with human hosts. Indicated percentage of strains carrying the respective sRNA, presence of sRNA was considered when observed over 60% coverage and 70 % nucleotide identity.

**Supplementary Figure S2. Growth of S. copri Wt and Δ*htcs*$^{Inu}$ with inulin as the sole carbon source. A.** Growth curve in minimal media supplemented with Inulin at 5 g/l as sole carbon source. Strains were grown in a 96-well plate under anaerobic conditions for up to 60h. Optical density was recorded every hour. The mean of three replicates is represented. **B.** Northern blot detection of SrcF in *S. copri* Wt grown 0.5x BHI supplemented with indicated monosaccharides. Samples were collected upon 20 h incubation. 5S rRNA was detected as a loading control.

**Supplementary Figure S3. Alignment of HTCS regulators for fructan degradation.** Alignment of HTCS regulators for Inulin or levan dedicated PULs. *Segatella copri* HTCS$^{Inu}$, *Bacteroides thetaiotaomicron* HTCS$^{Lev}$ VPI-5482, *Bacteroides uniformis*, *Bacteroides caccae* ATCC 43185, *Bacteroides ovatus* ATCC 8483, *Bacteroides caecimuris* I48. Domains of the HTCS are indicated with arrows. Residues described to interact directly with fructose in *B. thetaiotaomicron* HTCS$^{Lev}$ are indicated by black squares.

**Supplementary Figure S4. Polysaccharide utilization loci (PULs) expression in the mouse gut.** Heat map depicting transcriptional expression of annotated PULs in *S. copri* HDD04 at Stationary phase (n=3), from OMM12 mice cecum (n=2) and OMM12 mice colon (n=2) compared to *S. copri* grown to mid-exponential phase (n=3). $Log_2$ fold change in expression determined by DESeq2 is indicated. Loci and assigned PUL are indicated.

**Supplementary Figure S5. Metagenome analysis of *S. copri* colonized individuals with high and low SrcF expression. A.** Correlation analysis of SrcF expression in transcript per kilobase million (TPM) and relative abundance of *S. copri*. **B.** Alpha diversity analysis of high and low SrcF expressing microbiomes, Shannon (left panel) and observed diversity (right panel). **C.** Principal component analysis of Beta-diversity of high and low SrcF expressing microbiomes. **D.** Principal component analysis of cazymes composition of high and low SrcF expressing microbiomes.

**Supplementary Figure S6. Model of SrcF regulation and role in *S. copri*.** Graphical representation of regulatory pathways described. Degradation of fructans by *S. copri* or cohabitating commensals to modulate SrcF sRNA levels to regulate *S. copri* gene expression essential for successful colonization.

#### Supplementary Table legends:

**Supplementary Table S1. A.** *Segatella copri* HDD04 transcriptional start sites. Includes TSSs from growth in vitro. **B.** from OMM12 colonized mice and **C.** predicted terminators. **D.** *Segatella copri* DSM18205 transcriptional start sites. Includes TSSs from growth in vitro and **E.** predicted terminators. **F.** *Segatella copri* HDD04 small RNA annotations.

**Supplementary Table S2.** Expression of *S. copri* sRNAs in human donors Mikrodivers. **A.** *S. copri* HDD04 donor. **B.** *S. copri* HDA03 donor. **C.** *S. copri* HDB01 donor. **D.** *S. copri* HDC01 donor. **E.** Expression of *S. copri* sRNAs in human metatranscriptomes from NCBI-SRA BioProject: PRJNA354235.

**Supplementary Table S3. A.** 16S rRNA amplicon sequencing of *S. copri* colonized OMM12 mice colonization for dRNA-seq. **B.** and **C.** 16S rRNA amplicon sequencing of *S. copri*-colonized OMM12 mice of SrcF and HtcsInu competitions. **D.** Total RNA-seq of *S. copri* HDD04 ΔSrcF vs Wt at stationary phase. **E.** Total RNA-seq of *S. copri* HDD04 Wt in vitro vs colon of *S. copri*-OMM12 colonized mice. **F.** Total RNA-seq of *S. copri* HDD04 Wt in vitro vs cecum of *S. copri*-OMM12 colonized mice.

**Supplementary Table S4.** Cazymes abundance in metagenomes with high and low SrcF expression.

**Supplementary Table S5.** Strains, plasmids, and oligonucleotides.

**Supplementary Table S6. A.** MAG identifiers and respective classified *Segatella* species used for sRNA conservation analysis. **B.** Publicly available paired metatranscriptomic and metagenomic samples downloaded from NCBI-SRA BioProject: PRJNA354235. Metagenomes were excluded if the paired metatranscriptomes were not available or SrcF (TPM) was zero.

### Supplementary Figure S1

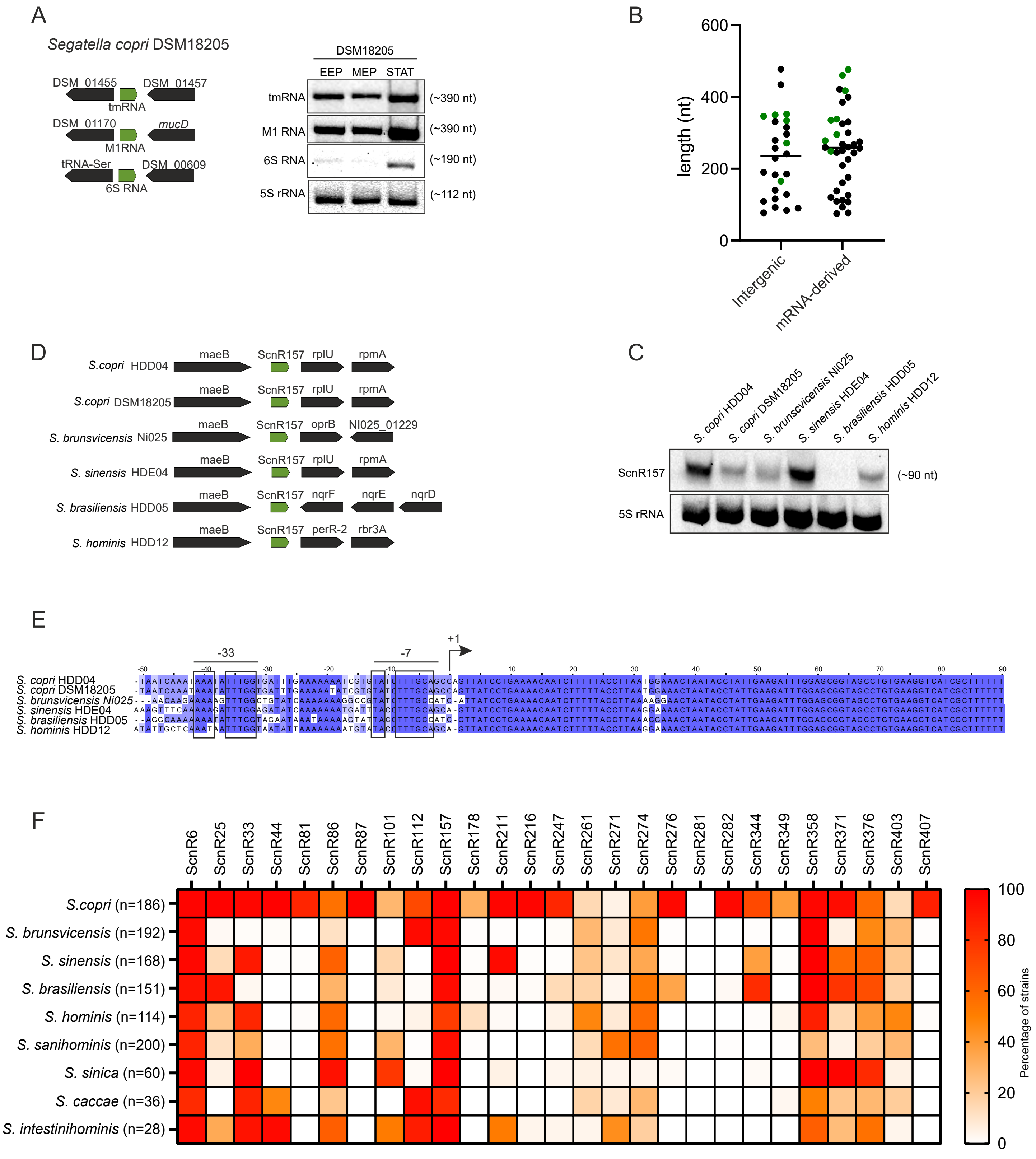

Supplementary Figure S2

A

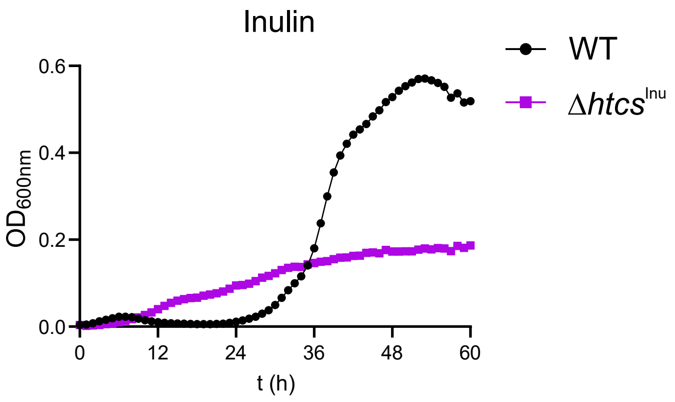


B

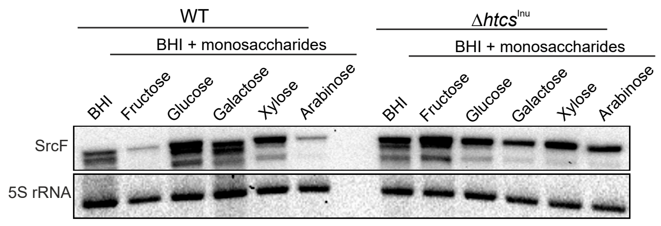

### Supplementary Figure S3

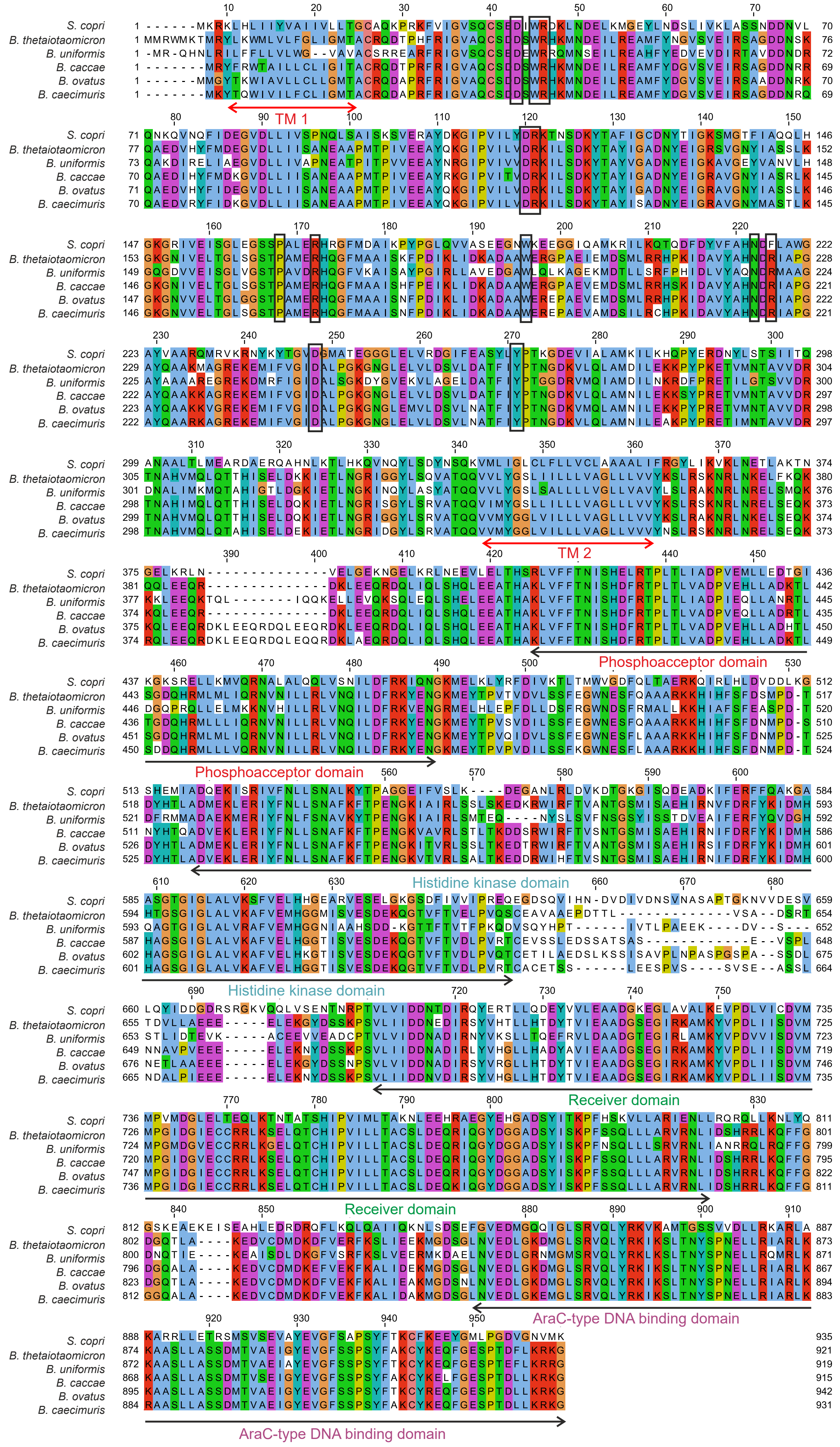

Supplementary Figure S4

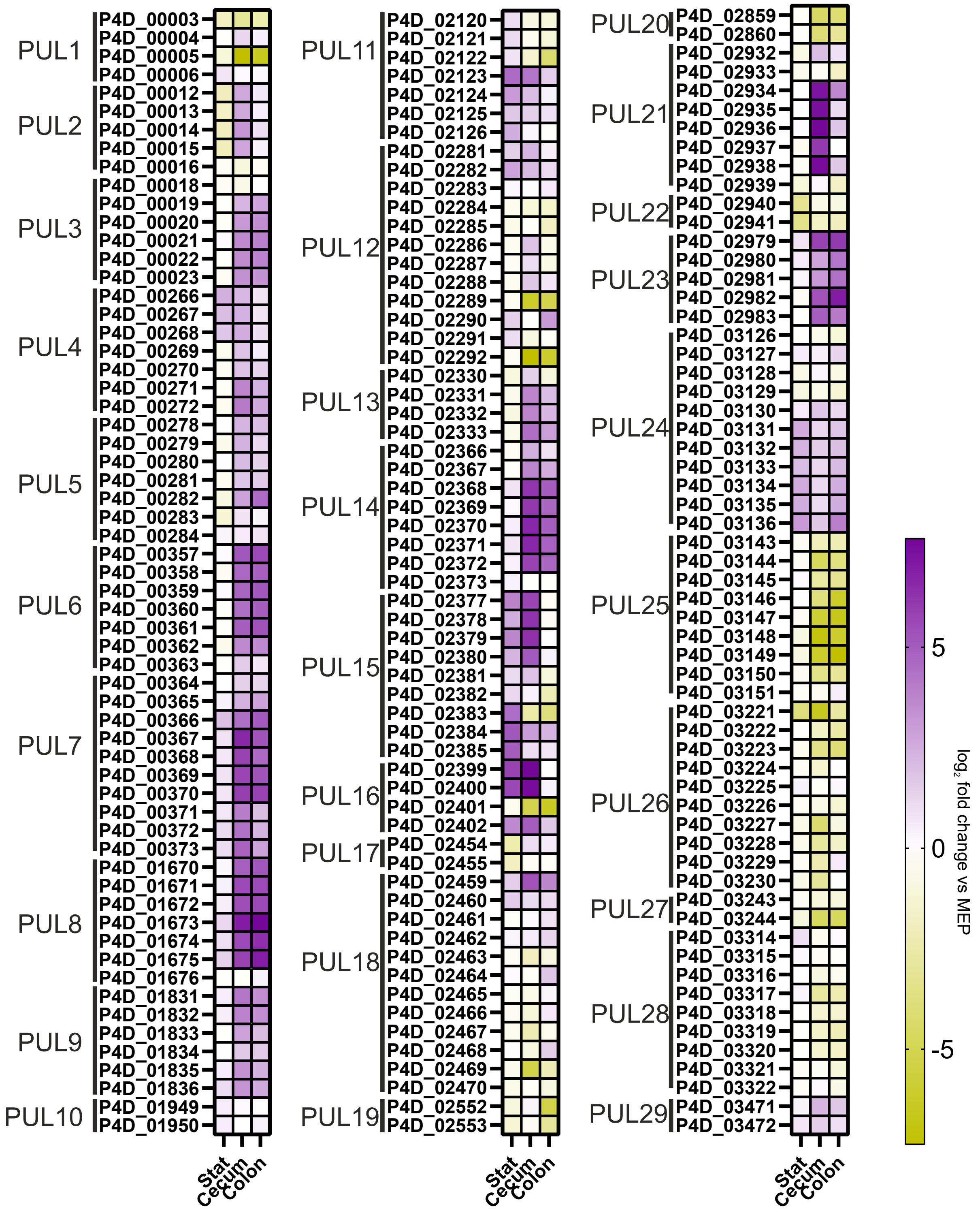

### Supplementary Figure S5

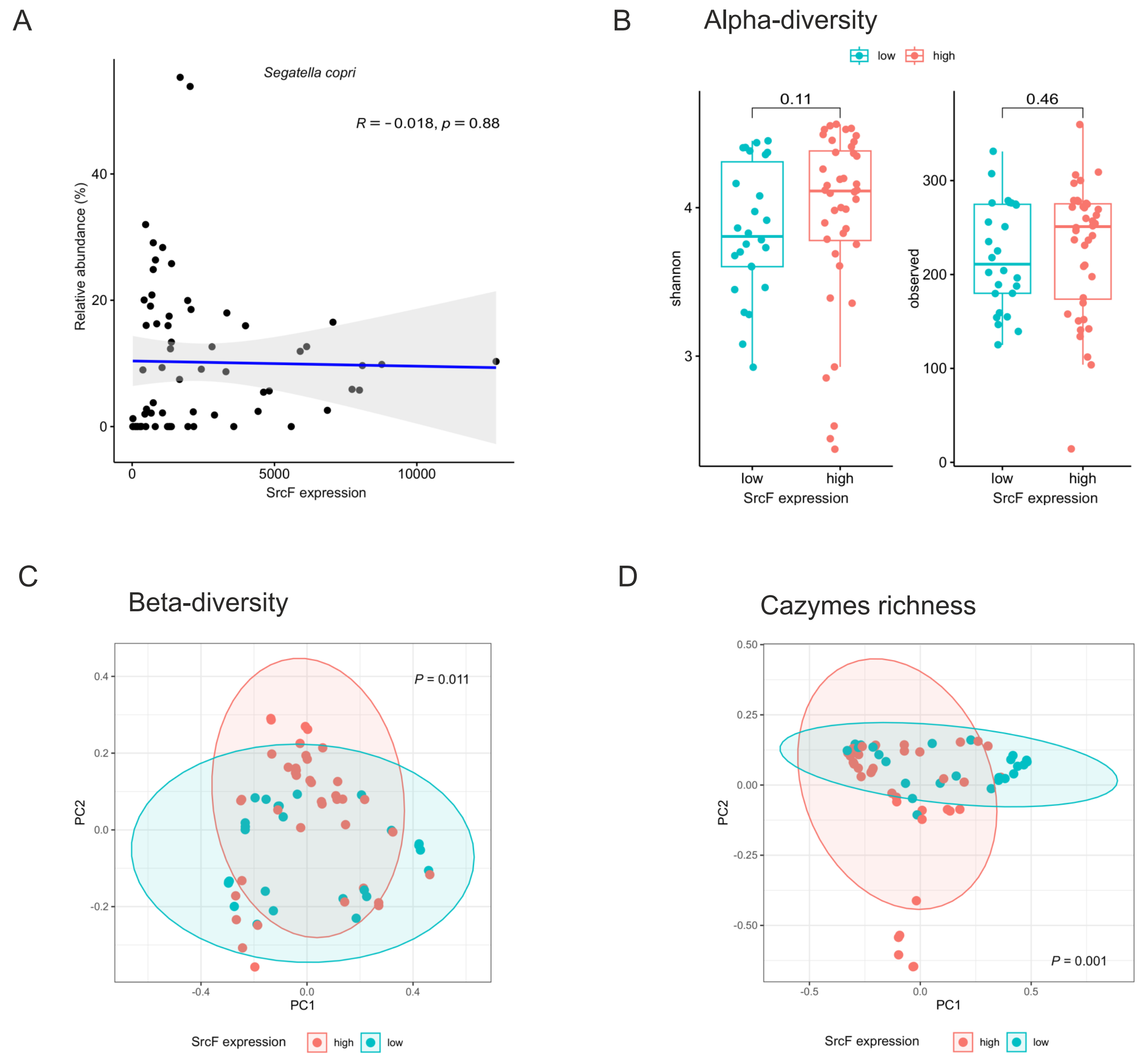

Supplementary Figure S6

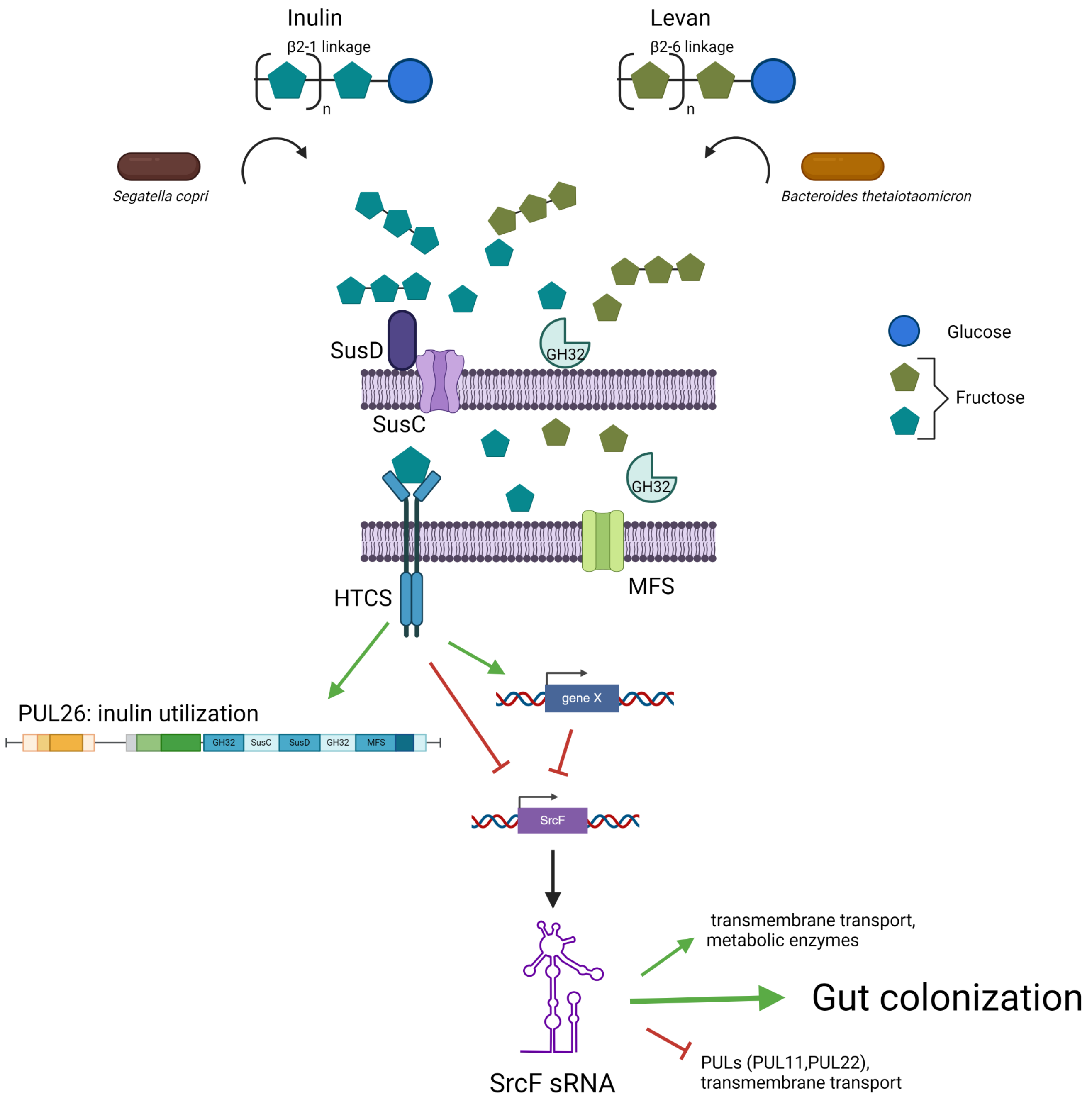
